# The machine-learning classifier ALLCatchR2 identifies 20 T-ALL subtypes across cohorts and age groups

**DOI:** 10.64898/2026.01.01.697268

**Authors:** Thomas Beder, Nadine Wolgast, Wencke Walter, Sonja Bendig, Alina M. Hartmann, Malwine J. Barz, Marketa Zaliova, Elisa Reitzel, David Baden, Stefan Schwartz, Nicola Gökbuget, Lennart Kester, Jan Trka, Claudia Haferlach, Monika Brüggemann, Claudia D. Baldus, Martin Neumann, Lorenz Bastian

## Abstract

T-cell acute lymphoblastic leukemia (T-ALL) comprises molecularly diverse subtypes, but robust cross-cohort validations and operational gene-expression definitions are lacking. To establish a gene-expression-anchored framework for T-ALL subtyping, we aggregated 2,314 transcriptomes (15 cohorts, age: 0.8 to 90.8 years). An extended unsupervised approach defined 17 main clusters and 3 subclusters in samples with high blast fractions. Supervised analyses added an overarching immature T-ALL (ETP-like) definition and resolved the LMO2 *γδ-like* subtype. All clusters contained samples from at least two cohorts. Characteristic genomic driver enrichments were consistent across cohorts, while gene expression clusters did not correspond exclusively to single driver events but also reflected developmental origins. A machine learning classifier based on ALLCatchR, our B-ALL classifier, identified these 20 transcriptomic subtypes and the immature T-ALL (ETP-like) signature with 0.995-1.0 accuracy in a validation set (n=203). Testing the classifier on a second hold-out data set (n=265 samples) showed that 92.7% of predictions matched with corresponding driver alterations. Across all samples, 83.2% of cases received high-confidence predictions, 7.3% candidate predictions, and 9.5% remained unclassified, largely because of low blast fractions. We identified a novel gene expression cluster markedly enriched (*P*<0.001) for clonal hematopoiesis mutations (*IDH2 R140Q, DNMT3A*) and a stem-/progenitor cell-like gene expression. This novel ‘clonal hematopoiesis-related’ T-ALL subtype was observed in six cohorts representing 8.9% of adults and 39.5% of patients aged >50 years. We advanced ALLCatchR, as a free R package that now enables B-/T-lineage separation, gene-expression subtyping, blast estimation, and developmental annotation to harmonize T-ALL classification across studies and clinical contexts.

**Key Points:**

1. Established using 2,314 T-ALL transcriptomes, ALLCatchR2 assigns 20 RNA-Seq subtypes and their developmental underpinnings across ages.
2. Clonal hematopoiesis-related T-ALL (*DNMT3A*, *IDH2 R140Q*) defines an immature gene-expression cluster in ∼40% of patients aged >50 years.

## Introduction

T-cell acute lymphoblastic leukemia (T-ALL) remains a therapeutic challenge, largely because of the absence of established immunotherapies that have recently improved outcomes in B-ALL and the limited availability of alternative targeted interventions. Molecular ALL subtypes reflect signaling trajectories downstream of groups or individual oncogenic drivers. They provide insights into the functional underpinnings of leukemogenesis - and potentially into targetable dependencies. Several studies have combined molecular genetics and gene-expression profiling to characterize the molecular landscape of T-ALL.^1–7^ Resulting subtype definitions such as TAL/LMO, TLX1, TLX3 or HOXA have been repeatedly identified in different cohorts. However, limited cohort sizes, a focus on specific age groups, and technical heterogeneity have impeded the establishment of robust cross-cohort subtype definitions. This is reflected in current disease classifications, which recognize only the early T precursor lymphoblastic leukemia (ETP-ALL; WHO-HAEM5^8^) or ETP-ALL with or without BCL11B rearrangement (ICC^9^) as distinct T-ALL subtypes supplemented by eight provisional entities in the ICC.^9^ This contrasts sharply with the 15 subtype definitions proposed by a recent landmark analysis.^10^

To date, the only available tool for systematic gene expression-based subtype allocation of individual T-ALL samples is limited to only nine subtype definitions.^11^ To establish robust and systematically validated T-ALL subtypes, we integrated RNA-Seq data from 2,314 T-ALL patients across 15 cohorts (age: 0.8–90.8 years) and identified 20 distinct transcriptional clusters, and an overarching immature ‘ETP-like’ group. This analysis independently confirmed rare subtypes and revealed a novel subtype enriched for mutations associated with clonal hematopoiesis.

## Methods

To define a transcriptomic reference of T-ALL and train the subtype machine learning classifier ALLCatchR2, we aggregated RNA-Seq count data from 15 cohorts^12–17,5,18–20,6,7,10,21^ comprising 2,314 samples (**Supplementary Table 1**). Gene-expression subtype discovery, ALLCatchR2 training and validation was performed using 2,049 samples from 14 cohorts, with independent testing conducted on 265 TARGET T-ALL^4^ samples. For samples from public resources, annotations including age, sex, blast counts and T-ALL subtype were obtained from the corresponding publications. Raw RNA-Seq FASTQ data from 307 in-house and 277 public T-ALL samples from 7 cohorts were analyzed (summary: **Supplementary Table 2**). Fusion calling and GATK variant calling was performed using IntegrateALL.^22^ For establishment of T-ALL gene-expression subtype definitions, gene-expression count data from 2,049 samples were harmonized across cohorts based on 17,361 genes detected in all data sets, and batch correction was performed using ComBat-seq.^23^ Corrected counts were normalized using log₂(count+1) transformation, followed by z-score scaling across samples and used for systematic unsupervised identification of gene expression clusters and annotation based on previous subtype assignments and driver calls obtained from the contributing cohorts. In a second step, the machine-learning classifier for T-ALL subtype allocations was developed using the original count data without cross-cohort count normalization. Counts were normalized within each sample separately to avoid data leakage. All methods are described in detail in the Results section and **Supplementary Materials**.

## Results

### Establishment of a representative age-spanning T-ALL gene expression reference cohort

A transcriptomic reference of T-ALL was established by aggregating in-house (n=443) and publicly available (n=1,606) RNA-Seq data sets from 14 cohorts (**Supplementary Table 2**). Patient age ranged from 0.8 to 90.8 years (66.0% pediatric / 34.0% adult). After batch correction, 2,049 T-ALL samples clustered independently of the contributing cohort (**Figure 1A**). Low-blast-count samples tended to group toward the center of the UMAP space (**Figure 1B**). Because low blast counts interfered with unsupervised identification of gene-expression subtypes, and measured blast counts were available for only 1,317 samples (64.3%), we trained a machine-learning regression model for T-ALL blast fractions. Training was performed on n=1008 samples^10^ achieving a correlation of measured to predicted blast counts of 0.725 (**Supplementary Figure 1A/B**). Testing on two hold-out cohorts achieved correlations of 0.532 and 0.696 (**Supplementary Figure 1C**).

**Figure 1.**
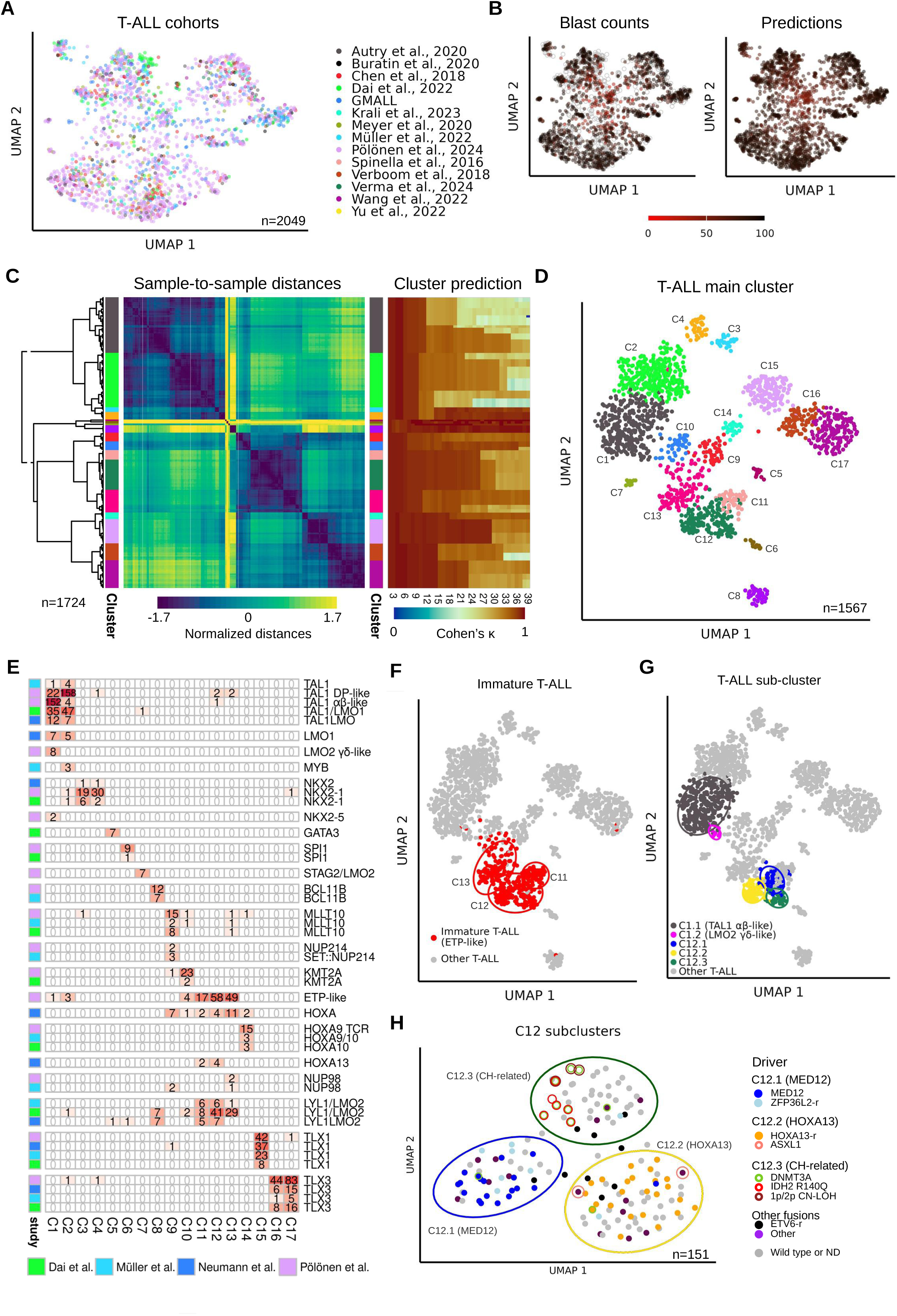
Systematic analysis of T-ALL samples revealed 20 T-ALL gene-expression subtypes. (**A**) UMAP plot showing batch-corrected count data of 2,049 T-ALL samples from 14 cohorts. The analysis was based on the 400 most variably expressed genes. (**B**) Same plot as in (A) but samples colored according to blast count and blast count predictions. Low blast count samples grouped towards the center of the UMAP plot. (**C**) Hierarchical clustering of 1,724 T-ALL samples with predicted blast counts > 70% using integrated sample-to-sample distances. The distances were averaged across 24 UMAP analyses with systematic variation of the parameters “min_dist” and “n_neighbors” and are shown on the left heatmap. The dendrogram was progressively split and at each split the integrity of the resulting clusters was determined by machine-learning (right heatmap). If Cohens’κ dropped below 0.8 the cluster was considered unstable, and the previous clustering level before the split chosen as the last stable group. (**D**) UMAP plot based on 1,330 lasso genes shows distinct clustering of 1,567 core representative samples of the 17 main gene-expression cluster identified by the unsupervised analysis. (**E**) Comparison of gene expression cluster to defined subtype definitions from the four largest cohorts. The number of samples for each cluster and the corresponding subtype definitions are indicated. (**F**) Same UMAP plot as in (D) but predicted immature T-ALL (ETP-like) samples are highlighted. (**G**) Subdivision of C1 into clusters C1.1 (TAL1 αβ-like) and C1.2 (LMO2 γδ-like), and C12 into three subgroups are highlighted. (**H**) C12 subclusters group distinctly in the UMAP analysis using 322 LASSO genes and are enriched for specific drivers indicating C12.3 as a novel subtype characterized by SNVs in the clonal hematopoiesis (CH) associated genes *DNMT3A* and *IDH2 R140Q*. The ellipses in the plot are based on the 95% confidence region.

T-ALL blast count models were implemented in our ALLCatchR^24^ framework which now allows blast-count predictions in B-ALL and T-ALL (**Supplementary Figure 2A**). To direct the classifier to the correct lineage, we used n=339 T-ALL and n=884 B-ALL samples from two cohorts to train a lineage classifier that achieved 0.98 accuracy. Predictions for T-lineage immunophenotype had a sensitivity of 0.98 and specificity of 0.99 in the hold-out data (**Supplementary Figure 2B**).

### Unsupervised identification of 17 T-ALL main gene expression clusters

To identify robust gene-expression clusters across cohorts, we selected high-blast-count samples (predicted infiltration >70%; n=1,724) for unsupervised analysis. As we have previously described,^25^ the 400 most variably expressed genes (**Supplementary Table 3**) were used to generate 24 UMAP spaces by systematic parameter variation (**Supplementary Figure 3A**). Sample-to-sample distances were normalized, averaged across these spaces and subjected to hierarchical clustering (**Figure 1C**). Cluster robustness was determined by iteratively splitting sub-clusters and testing the ability of a machine-learning classifier to reproduce the corresponding cluster. Once Cohen’s κ for predictability dropped below 0.8, the next node above was considered the last stable entity. This approach identified 17 major T-ALL gene expression clusters predictable with a median sensitivity of 0.929 (±0.073), specificity of 0.999 (±0.002) and κ of 0.939 (±0.047), demonstrating robustness of definitions obtained from unsupervised analysis (**Supplementary Figure 3B**). Of 1,724 samples, 1,605 (93.1%) were assigned to a unique cluster by the machine learning classifier, whereas 119 samples remained unclassified (**Supplementary Figure 3C**). Unclassified samples often localized between clusters (**Supplementary Figure 3D**). From 1,605 samples with subtype assignment, 1,567 (97.6%) were correctly allocated to their initial gene expression cluster (**Figure 1D**). We used subsequently this cohort as core representatives of their corresponding subtype for ALLCatchR2 training and validation. Importantly, no exclusively age-driven subclusters emerged from the unsupervised cluster identification, arguing against age as the dominant determinant of cluster definitions.

### Cross-cohort definition of gene expression subtypes

Analysis of subtype definitions in the four largest contributing cohorts^5–7,10^ showed that most of the clusters mapped clearly to pre-defined entities (**Figure 1E, Supplementary Table 4**), including clusters *C1 (TAL1 αβ-like, LMO2 γδ-like), C2 (TAL1 DP-like)*, *C6 (SPI1)*, *C7 (STAG2/LMO2)*, *C8 (BCL11B)*, *C9 (MLLT10, NUP214), C10 (KMT2A)*, *C14 (HOXA9/10 TCR)* and *C15 (TLX1)*.

Cluster C5 included 10 samples with a highly distinct gene expression profile (**Figure 1D**) and was initially defined by *GATA3* single nucleotide variants and a corresponding ‘GATA3’ subtype (7/7) exclusively described in one contributing cohort^5^ (**Figure 1E**). Recent analyses described a T-ALL / MPAL subtype characterized by a t(14;16)(q32;q24) translocation, which places the *FOXF1* / *FENDRR* locus under the regulatory control of the *BCL11B* enhancer^26,27^. Common secondary events of this FOXF1/FENDRR subtype included loss-of-function *GATA3* mutations in all cases. Single-sample GSEA using top differentially expressed genes^26^, confirmed that cluster C5 corresponded to the novel FOXF1/FENDRR subtype (**Supplementary Figure 4A**). In addition, FOXF1 and FENDRR expression was consistently high in C5 cases (**Supplementary Figure 4B**). Published FOXF1/FENDRR immunophenotypes showed a B-/T-lineage ambiguity with co-expression of T-cell markers cytoplasmatic CD3, CD7 and CD19, CD79A, CD10. Nevertheless, the most frequent diagnostic immunophenotype had been ETP-ALL and cluster C5 samples grouped mostly with T-ALL in our B- vs.T-ALL UMAP analysis. Therefore, we kept this subtype in our T-ALL classifier as C5 (FOXF1/FENDRR).

Cluster C3 and C4 represented two NKX2-1-driven clusters: *C3 (NKX2-1 other)* was enriched for chromosome 14 chromothripsis (6/6, *P*<0.01, **Supplementary Figure 5**, **Supplementary Table 5**), *NKX2-1* intergenic loss (5/6, *P*<0.05) and *MYB::TCR* (5/12, *P*<0.001), whereas *C4 (NKX2-1::TCR)* was enriched for *NKX2-1::TCR* gene fusions (27/33, *P*<0.001). Similarly, *C16 (TLX3 DP-like)* and *C17 (TLX3 immature)* were both enriched for TLX3 subtype and characterized by *TLX3::BCL11B*-enhancer fusions (112/117). However, *C17 (TLX3 immature)* was enriched for ETP/nearETP immunophenotype (*P*<0.001) and also showed additionally enrichment for *NUP214::ABL1* (27/43, *P*<0.001) and other fusions (3/4 *TLX3::CDK6* and 7/12 *TLX3::TCR*). Both, *NKX2-1* and *TLX3*-sub-clusters, had been described previously in one pediatric cohort^10^ and the reproducibility of these and the other subtypes in independent cohorts confirmed their existence also in adults.

Despite the large overlap between established subtypes and gene expression clusters, discrepancies were observed as well. These can be exemplified with seven samples in the well-defined NKX2-1, TLX1 and TLX3 subtypes in the original publications that did not group within the corresponding gene expression cluster. Based on the expression of 1,330 genes, these samples grouped clearly to gene expression clusters, not to their defined subtypes (**Supplementary Figure 6A**). Moreover, the marker expression levels of *NKX2-1* and *TLX3* did not match the subtype definitions in the original publications but the gene expression cluster assignments (**Supplementary Figure 6B**). Beyond that, the genomically defined subtypes NKX2-5 (n=2), MYB (n=3) and NUP98 (n=5) did not form individual gene expression clusters, potentially due to the small number of samples, or less distinct expression signatures. Instead, these cases were included within main cluster definitions (**Figure 1E**) illustrating the limitations of gene expression based classification of very rare genetic subtypes.

### Definition of driver overarching immature T-ALL (ETP-like)

C11-C13 were enriched for the recently described ETP-like subtype^10^ (124/132, *P<0.001,* **Figure 1E**). To establish an operationalizable definition of this driver overarching entity in our cross-cohort data set, we used the differentially expressed genes between previously defined^10^ ETP-like (n=132) and non-ETP-like (n=665) cases (**Supplementary Figure 7A**). Machine-learning classification for the resulting two clusters achieved an accuracy of 0.987 (**Supplementary Table 6**) with marker gene expression confirming a higher expression of stem cell and early myeloid markers in ETP-like cases including CD34, CD117 and CD33 (*P*<0.001) and lower expression T-cell differentiation of markers such as CD2, CD1a, CD4 and CD8 (*P*<0.001) consistent with previous immunophenotypic definitions (**Supplementary Figure 7B**). To distinguish the gene expression definition from immunophenotypically defined ETP T-ALL, we termed this cluster overarching group ‘immature T-ALL (ETP-like)’.

Despite this unifying immature T-ALL (ETP-like) definition, most of these samples (n=311/324) belonged to three individual clusters from the unsupervised analysis: C11, C12 and C13 (**Figure 1F**). Cluster *C11 (immature ZFP36L2)* was enriched for *ZFP36L2* gene fusions (9/15, ZFP36L2::TCR, *P<0.001; 2/6, ZFP36L2::SFPQ, P<0.05*) while *C12 (immature MED12, HOXA13, ZFP36L2)* was enriched for *MED12* mutations (19/36, *P<0.001)*, *HOXA13* aberrations (25/25*, P<0.001*), *ZFP36L2::TCR* (4/15*, P<0.05*) and *ZFP36L2::SFPQ* (4*/6, P<0.05*) gene fusions. *C13 (immature NUP214, MLLT10, KMT2A)* contained drivers also found in C9 and C10 indicating that C13 grouped samples according to an overarching immaturity phenotype beyond these specific drivers (**Supplementary Figure 5**).

### Further subclassification of clusters C1 and C12

Cluster *C1 (TAL1 αβ-like, LMO2 γδ-like)* contained all n=8 LMO2 γδ-like samples that grouped distinctly from TAL1 αβ-like^10^ in UMAP space (**Figure 1G)**. To distinguish the small but clinically relevant LMO2 γδ-like subtype, we used top differentially expressed genes between cases annotated as TAL1 αβ-like (n=152) or LMO2 γδ-like (n=8) annotation in one cohort^10^ to cluster C1 cases of our multi-cohort data set, revealing a clear separation of both subtypes (**Supplementary Figure 7C**). Training a machine-learning model with randomized stratified cross-validation achieved an accuracy of 0.984 for separating these two subtypes (**Supplementary Table 7**), which otherwise remained challenging due to the scarcity of LMO2 γδ-like samples and shared driver alterations, leaving the developmental stage of arrest as the primary determinant of the gene expression phenotype^10^.

UMAP analysis showed that C12 (immature MED12, HOXA13, ZFP36L2) samples further separated within this cluster according to their corresponding drivers (**Supplementary Figure 8A**). To assess the robustness of this observation, we applied the iterative clustering approach used to define the 17 main clusters to the 151 samples of cluster C12 only (**Supplementary Figure 8B**, **Supplementary Table 8**).

The resulting *C12.1 (MED12)* subcluster comprised almost all cases with reported *MED12* mutations (18/19, *P*<0.001) but also some *ZFP36L2* gene fusions (**Figure 1H**). In comparison with C11 (*immature ZFP36L2*) samples, which frequently harbored *ZFP36L2* rearrangements to the TCR locus, samples in C12.1 were enriched for other *ZFP36L2* translocation partners (e.g., *ZFP36L2::SFPQ* (*P*<0.05)), and 131 genes were differentially expressed between *ZFP36L2* aberrant samples of both clusters. The *C12.2 (HOXA13)* subcluster contained all samples with *HOXA13* aberrations (25/25, *P*<0.001). *C12.3 (CH-related)* was enriched for clonal hematopoiesis (CH) related SNVs in *DNMT3A* (n=8/108, *P*<0.001) and *IDH2 R140Q* (n=6/6, *P*<0.001) and also contained 2/2 cases with CN-LOH 1p and 2/2 samples with CN-LOH 2p (**Figure 1H**). Interestingly, *ASXL1* mutations were not found in the *C12.3* but in the *C12.2 (HOXA13)* cluster. *ETV6-r* samples were found in all C12 sub-clusters (**Figure 1H**). DGEA comparing ETV6-r against all other C12 samples showed no differentially expressed genes (|L2FC| > 1.5 and FDR ≤ 0.05), indicating that additional factors drive expression differences within *ETV6-r*.

In summary, we identified 20 T-ALL gene-expression subtypes and a subtype-overarching definition of immature T-ALL (ETP-like) characterized by distinct gene expression profiles (**Supplementary Figure 9**).

### Development of a T-ALL subtype classification model in ALLCatchR2

Based on the definition of 20 T-ALL gene-expression subtypes and the overarching definition of immature T-ALL (ETP-like), we developed a systematic T-ALL subtype classifier. We split the 1,567 core representative samples from 14 cohorts (**Supplementary Table 2**) into training (n=1,364; 11 cohorts) and validation sets (n=203; 3 cohorts). The classifier comprises one module for predicting 20 T-ALL gene-expression clusters and another for identifying immature T-ALL (ETP-like). Initially, predictions were made for the 17 major gene-expression clusters. For samples assigned to cluster C1 or C12, subsequent predictions were performed to resolve the corresponding subclusters (**Figure 2A**). To find the most suitable type of machine-learning model, a total of ten were evaluated based on their ability to predict the 17 major gene expression subtypes in the validation samples (**Supplementary Figure 10**). For the final predictions, the scores from the svmLinear3, svmLinearWeights2, ranger, rf, and LogitBoost models were combined based on complementary benefits of this combination (more details: Supplementary Material and Methods). The combined prediction scores demonstrated high discriminative performance, effectively stratifying samples according to distinct gene-expression subtypes and immature T-ALL status. Cluster-specific score cutoffs (**Supplementary Table 9**) defined high-confidence, candidate, and unclassified calls on the training data (**Figure 2B**). ALLCatchR2 enabled hierarchical classification as exemplified in four samples in **Supplementary Figure 11A**.

**Figure 2.**
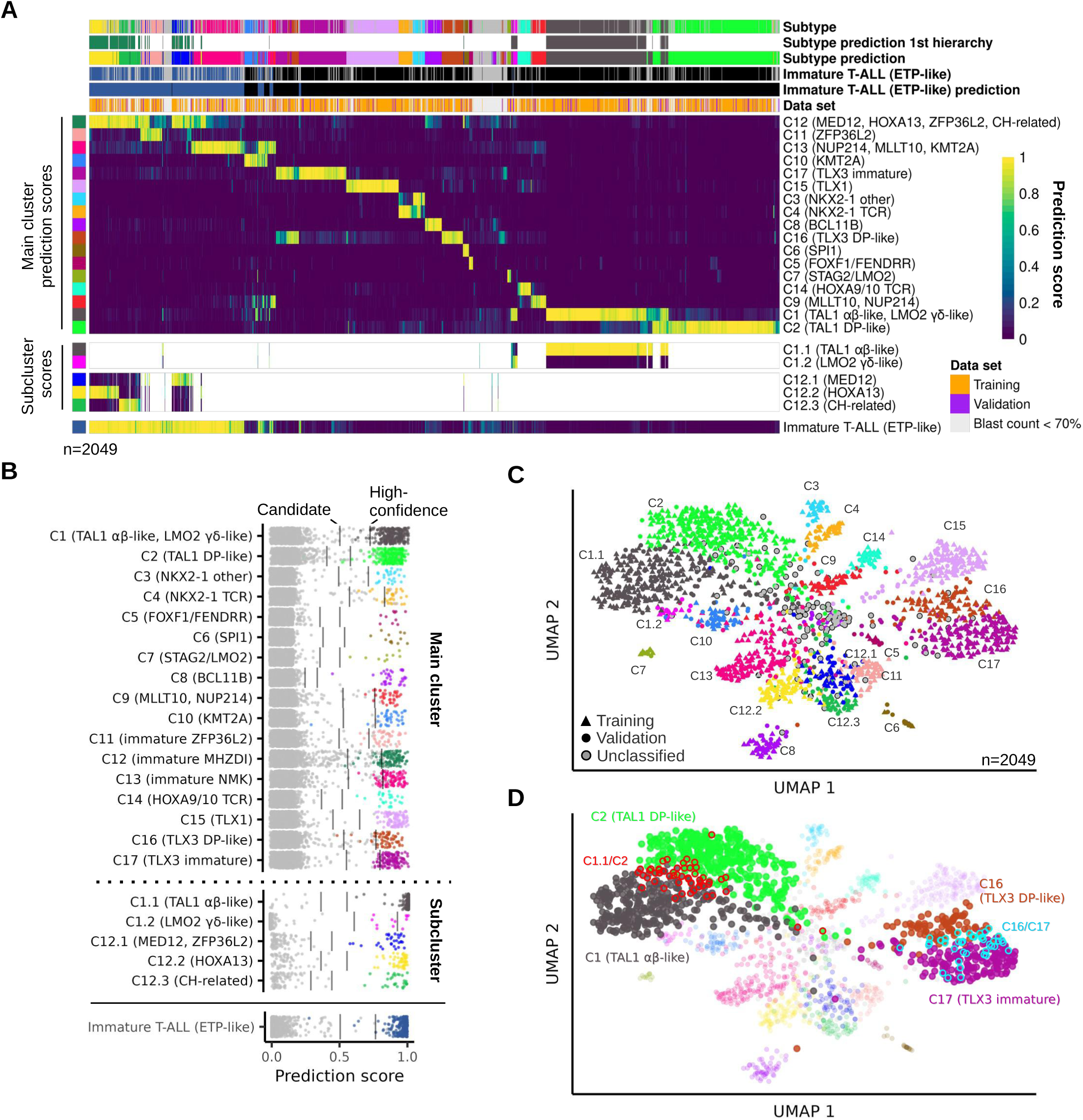
Implementation of ALLCatchR2 for prediction of 20 T-ALL subtypes and immature T-ALL (ETP-like). (**A**) Heatmap showing the prediction scores for gene expression-defined T-ALL subtypes of n=2,049 T-ALL samples. Core set samples (n=1,567) were assigned to subtypes and remaining samples considered “unassigned” and colored grey in the annotation rows. C1 and C12 predicted samples are indicated in “Subtype prediction 1st hierarchy” as these were further assigned into sub-entities. In addition, immature T-ALL (ETP-like) predictions from the second ALLCatchR2 module are shown. Unclassified samples are indicated by lightgrey color in the annotation rows. ALLCatchR2 was trained using 1,364 samples and validation was performed on 203 samples with subtype assignments that are indicated in the “Data set” annotation row. (**B**) Scatterplot showing prediction score distribution for T-ALL gene expression subtypes. For each subtype the corresponding samples, and high-confidence and candidate cutoffs are indicated. (**C**) UMAP plot based on 1,689 LASSO genes selected for the individual subtypes. Predicted subtypes are colored according to (A). (**D**) Same UMAP plot and samples with predictions for both clusters C1.1 (TAL1 αβ-like) and C2 (TAL1 DP-like), or C16 (TLX3 DP-like) and C17 (TLX3 immature) are indicated. Predictions of samples from the other clusters are transparent and shown for orientation.

Using the defined cutoff thresholds, the classifier achieved accuracies of 0.995 for main cluster prediction, 1.0 for subcluster prediction, and 0.995 for immature T-ALL (ETP-like) prediction across the 203 validation samples. Prediction scores for the 2/203 misclassified validation samples are shown in **Supplementary Figure 11B**. Sample A25 from cluster C12.1 (MED12) was predicted as C11 (immature ZFP36L2), and the immature T-ALL (ETP-like) sample GSM4087091 was misclassified as non-immature. We analyzed all 2,049 T-ALL samples from the 14 cohorts and 1,705 (83.2%) received high-confidence prediction scores, 149 (7.3%) were classified as candidate predictions, and 195 (9.5%) remained unclassified (**Figure 2A**). Unclassified samples exhibited significantly lower median blast counts (69%±13%) than samples with high-confidence (83%±9%, *P*<0.001) or candidate predictions (76%±12%, *P*<0.05) and grouped towards the center of the UMAP (**Figure 2C**; **Supplementary Figure 11C**).

T-ALL gene-expression subtypes also reflect the developmental stage of arrest and independent factors such as blast fraction. To account for this uncertainty, we set up a framework within the classifier that uses quantitative scores and enables assignment of samples to multiple subtypes when prediction uncertainty exists. Considering both high-confidence and candidate cutoffs, 1740/2049 (84.9%) samples received a single unambiguous subtype allocation, whereas 289/2049 (14.1%) and 20/2049 (1.0%) met the threshold for two or more subtype predictions, respectively (**Supplementary Table 10**). Only 102/2049 samples (5.0%) had two high-confidence subtype calls. The highest rate of overlapping subtype predictions affected cases in closely related clusters, i.e., co-classifications for C1.1 (TAL1 αβ-like) and C2 (TAL1 DP-like), C16 (TLX3 DP-like) and C17 (TLX3 immature) or the two *NKX2-1*-related clusters C3 and C4 (**Figure 2D**, **Supplementary Table 10**).

### Predicted subtypes are confirmed in independent hold-out samples

In the independent TARGET cohort^4^, ALLCatchR2 achieved high-confidence classifications in 94.0% of samples, candidate classifications in 4.5%, and only 1.5% of samples remained unclassified. UMAP projection using our subtype-defining gene-expression signature (**Supplementary Table 11**) revealed a well-defined cluster separation (**Figure 3A**). TARGET ALL samples had reported drivers from WGS/RNA-Seq integration^10^ in 94.0% of samples (**Supplementary Table 12**). To determine whether the enrichment patterns of genomic drivers in our ALLCatchR2 clusters were reproducible across cohorts, we aggregated all samples from 15 cohorts with available driver information (n=2,149/2,314, 92.9%). We split them into a ‘training’ set comprising 1,257 core samples and a hold-out test set for validation (n=892) including 198 core samples used for ALLCatchR2 validation, 436 samples that had initially been excluded from the core set because of low blast fractions and 258 TARGET samples. Significant associations with one of the ALLCatchR2 gene-expression subtypes were found for 43 drivers (**Figure 3B**, **Supplementary Table 13**). The highest correlations (ρ > 0.90) were observed for fusions such as *PICALM::MLLT10*, *HOX9/10::TCR*, *HOXA13::BCL11Benh, KMT2A::MLLT1*, *STMN1::SPI1*, *TLX1-r*, *TLX1NB-r* and *TLX3-r*, which were all highly specific for individual gene-expression subtypes (**Figure 3C**). Lower correlations were observed with rare drivers, e.g. *LOSS:NKX2-1* represented in five C3 and one C4 sample in training and in only one C4 sample of the validation cohort (**Supplementary Table 13**).

**Figure 3.**
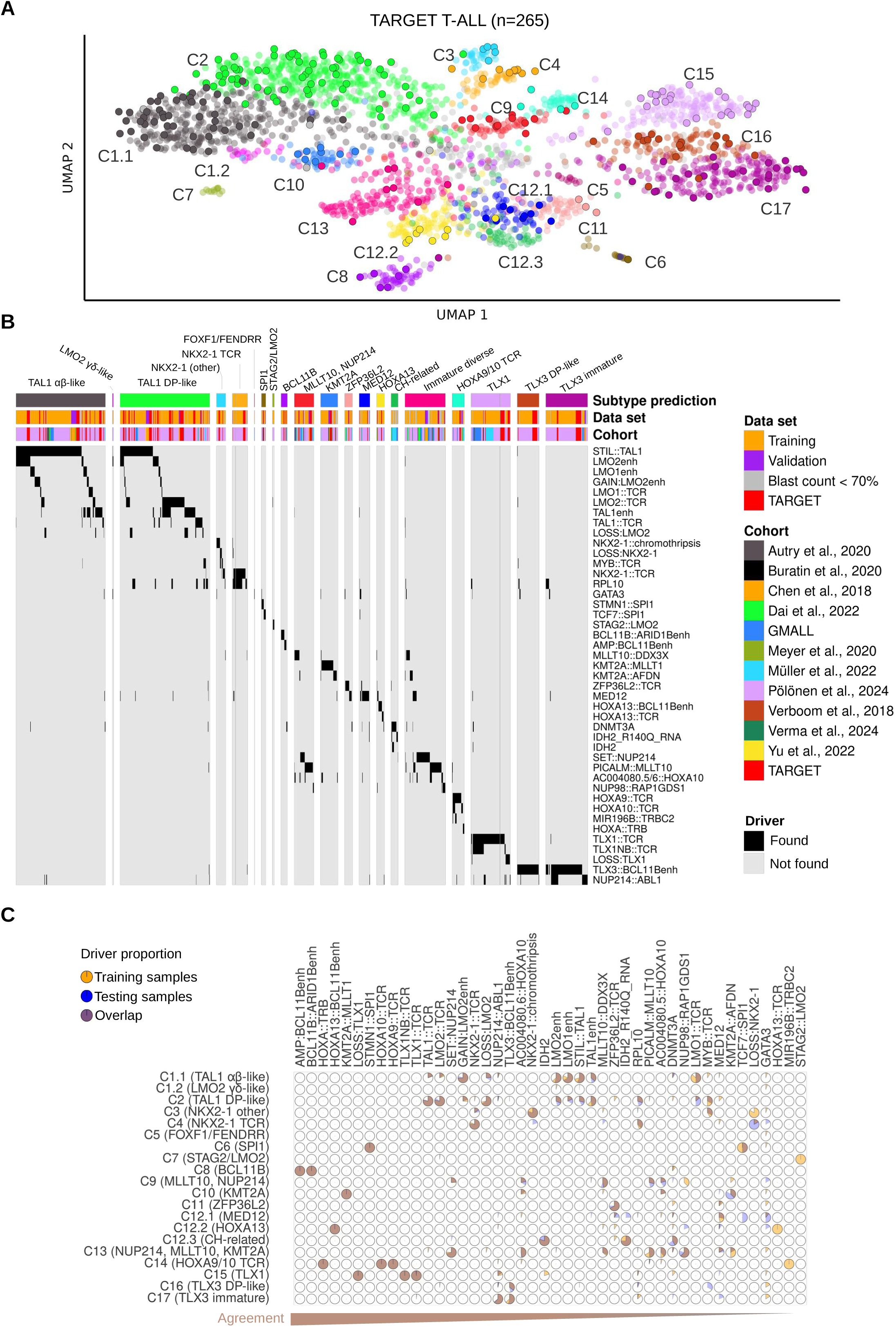
Evaluation of ALLCatchR2 T-ALL subtype predictions and driver presence. (**A**) Testing ALLCatchR2 on n=265 TARGET ALL samples not used for establishing the gene expression clusters or classifier training. TARGET samples were projected onto the UMAP in Figure 2C and are highlighted. (**B**) Heatmap showing 1,260 samples with drivers enriched in T-ALL gene expression subtypes. (**C**) Proportion of samples in the training and testing data for 43 significantly enriched drivers are shown. Drivers are sorted according to the correlation coefficient between training and testing samples.

### Oncogene expression patterns define T-ALL subtypes

T-ALL subtypes are characterized by oncogene overexpression downstream of different driver alterations. Comparison of driver oncogene expression across clusters confirmed upregulation of *TLX3*, *TLX1*, *NKX2-1* and *TAL1* in samples of the corresponding subtypes (**Figure 4A**). As reported, *TLX1* and *TLX3* subtype cases also showed high expression of *NKX2-1*, while isolated *NKX2-1* overexpression was specific to the *NKX2-1* subtype. Notably, both *TLX3* and *NKX2-1* subclusters had comparable expression levels of their corresponding oncogene. *HOXA* gene-expression patterns varied across subtypes: C9, C10, C12.3, and C14 showed increased expression of anterior *HOXA* genes (*HOXA5*, *HOXA9,* and *HOXA10*), whereas the posterior gene *HOXA13* was specifically higher expressed in C12.2, and C13 expressed both anterior and posterior *HOXA* genes. *MEF2C* and *LYL1* were highly expressed in immature T-ALL (ETP-like) but also in C8 (BCL11B) and C10 (KMT2A). To facilitate oncogene analysis in individual samples, we added an ALLCatchR2 module that maps oncogene expression against the reference cohort.

**Figure 4.**
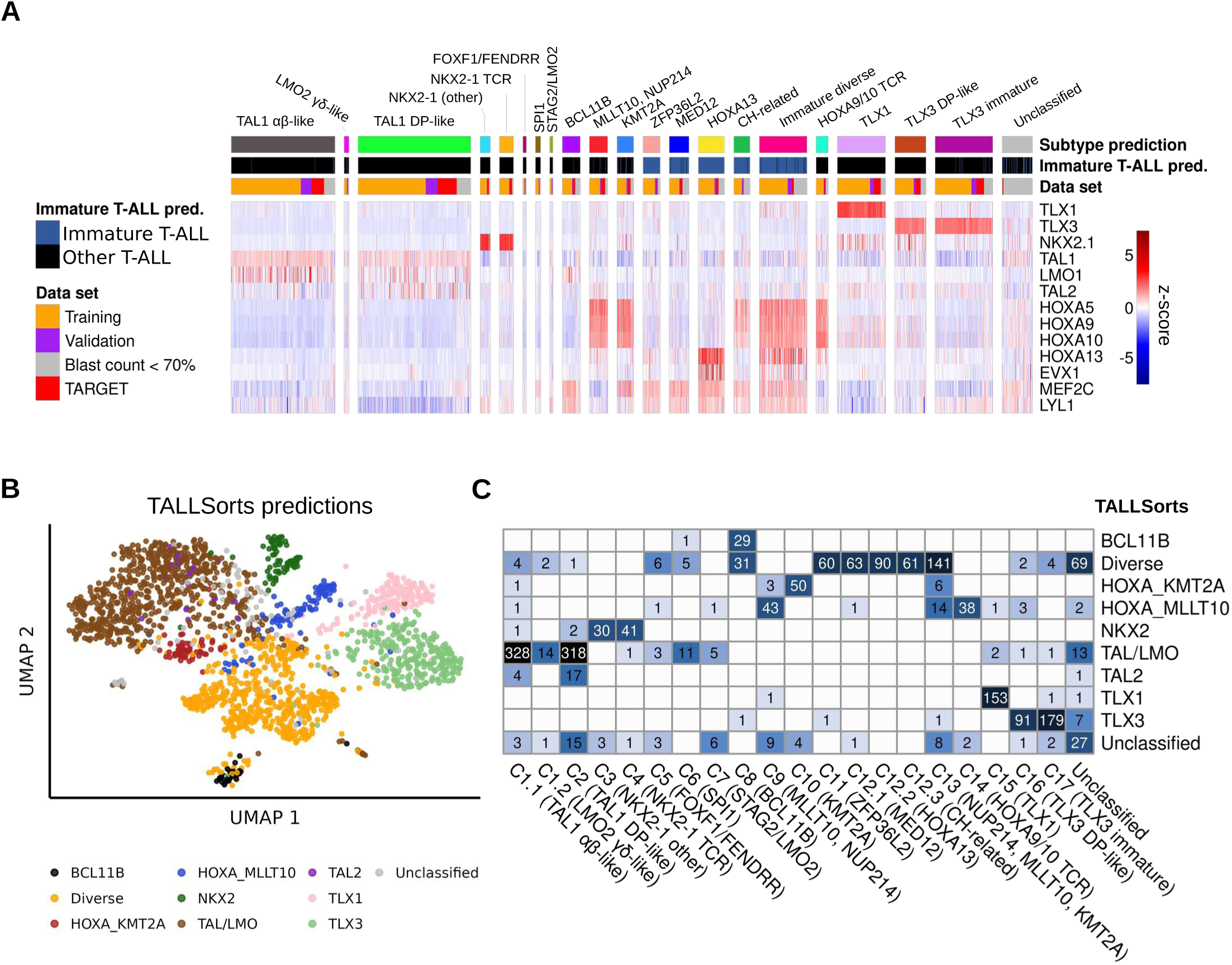
Marker gene expression and comparison of ALLCatchR2 to TALLSorts predictions. (**A**) Marker gene expression for each sample sorted by T-ALL subtypes. Expression of *TLX1*, *TLX3*, *NKX2-1* and *HOXA13* distinguished subtypes characterized by alterations in these genes. Training and testing samples are indicated. (**B**) Comparison of ALLCatchR2 and TALLSorts for gene expression-based T-ALL subtype classification. Same UMAP plot as in Figure 2C, but samples are colored according to TALLSorts predictions. (**C**) Heatmap showing the comparison between ALLCatchR2 and T-ALLSorts subtype predictions.

### ALLCatchR2 subtype definitions extend beyond the existing classifier

To our knowledge, TALLSorts^11^ is currently the only publicly available tool for T-ALL subtype classification based on gene-expression profiling only. TALLSorts assigned 95.8% of the 2,049 samples used to establish ALLCatchR2 models to nine broader subtypes (**Figure 4B**). These largely corresponded to several of the 20 subtypes defined in this study, indicating that ALLCatchR2 achieves higher prediction resolution (**Figure 4C**). Only the TLX1 and HOXA_KMT2A TALLSorts definitions were concordant with C15 (TLX1) and C10 (KMT2A). By contrast, NKX2 and TLX3 each corresponded to two ALLCatchR2 subtypes and HOXA_MLLT10 corresponded to C9 (MLLT10, NUP214) and C14 (HOXA9/10 TCR). Discrepancies were observed for C7 (STAG2/LMO2), for which 5/12 samples were predicted as TAL/LMO and 6/12 remained unclassified by TALLSorts. The TAL2 definition was enriched in C2 (TAL1 DP-like), but TAL2 samples did not form a distinct cluster (**Figure 4B**). C6 (SPI1) and C5 (FOXF1/FENDRR) were variably predicted as TAL/LMO or Diverse by TALLSorts (**Figure 4C**). The Diverse group included 31/61 C8 (BCL11B) samples and C11, all C12 subclusters and C13 also largely mapped to Diverse as well. Low-blast-count samples were also frequently predicted as Diverse (**Figure 4B**) and thus 69 samples in this group were not classified by ALLCatchR2.

### A novel T-ALL subtype defined by clonal hematopoiesis (CH)-associated gene mutations

The presence of *DNMT3A* (n=21/41 of all *DNTM3A* mutated cases in the cohort, *P*<0.001) *IDH2 R140Q* (n=13/16, *P*<0.001) and *TET2* (4/8 *P*<0.001) SNVs was characteristic of the novel C12.3 gene-expression cluster which we termed CH-related because of the frequent involvement of these genes in clonal hematopoiesis (**Figure 3C, Supplementary Figure 12**). To validate this distribution, we analyzed WGS data (n=121) from one of our own unselected adult cohorts (MLL). This identified *DNMT3A* and *IDH2 R140Q* mutations in 12/16 and 5/16 C12.3 cluster patients respectively, compared to 7/105 and 0/105 patients in the remaining cohort (**Figure 5A**; *P*<0.001 for both comparisons), confirming a strong enrichment of CH-driver mutations in this cluster. Patients in C12.3 (CH-related) had the highest median age (57.1±17 years; **Figure 5B**). In total, 61 samples were predicted to belong to the C12.3 (CH-related), accounting for 2.7% of all T-ALL cases, but 8.9% of adult cases (*P*<0.001) and 39.5% of patients older than 50 years (**Figure 5C**). In contrast, pediatric cases (<15 years of age) represented a broad diversity of clusters with a high proportion of TAL1-related subtypes (C1.1, C2), C7 (STAG2/LMO2), C6 (SPI1), C1.2 (LMO2 γδ-like), C16 (TLX3 DP-like), and both *NKX2-1* subtypes **(Supplementary Figure 13A**). Adolescents and young adults (AYA; 16-39 years of age) showed a composition partially overlapping with both pediatric and adult cohorts and were enriched for C5 (FOXF1/FENDRR). The adult representation of some clusters was mainly driven by AYA cases (e.g., C10, C3, C3, C4 or C5). Adult cases (>40 years of age) displayed a shift toward increased representation of C15 (TLX1), C11 (immature ZFP36L2), and C8 (BCL11B) which are rare in children under 10 years. Correlation analysis indicated a continuous shift from pediatric to adult age, reflecting the age-dependent selection of drivers (**Supplementary Figure 13B**).

**Figure 5.**
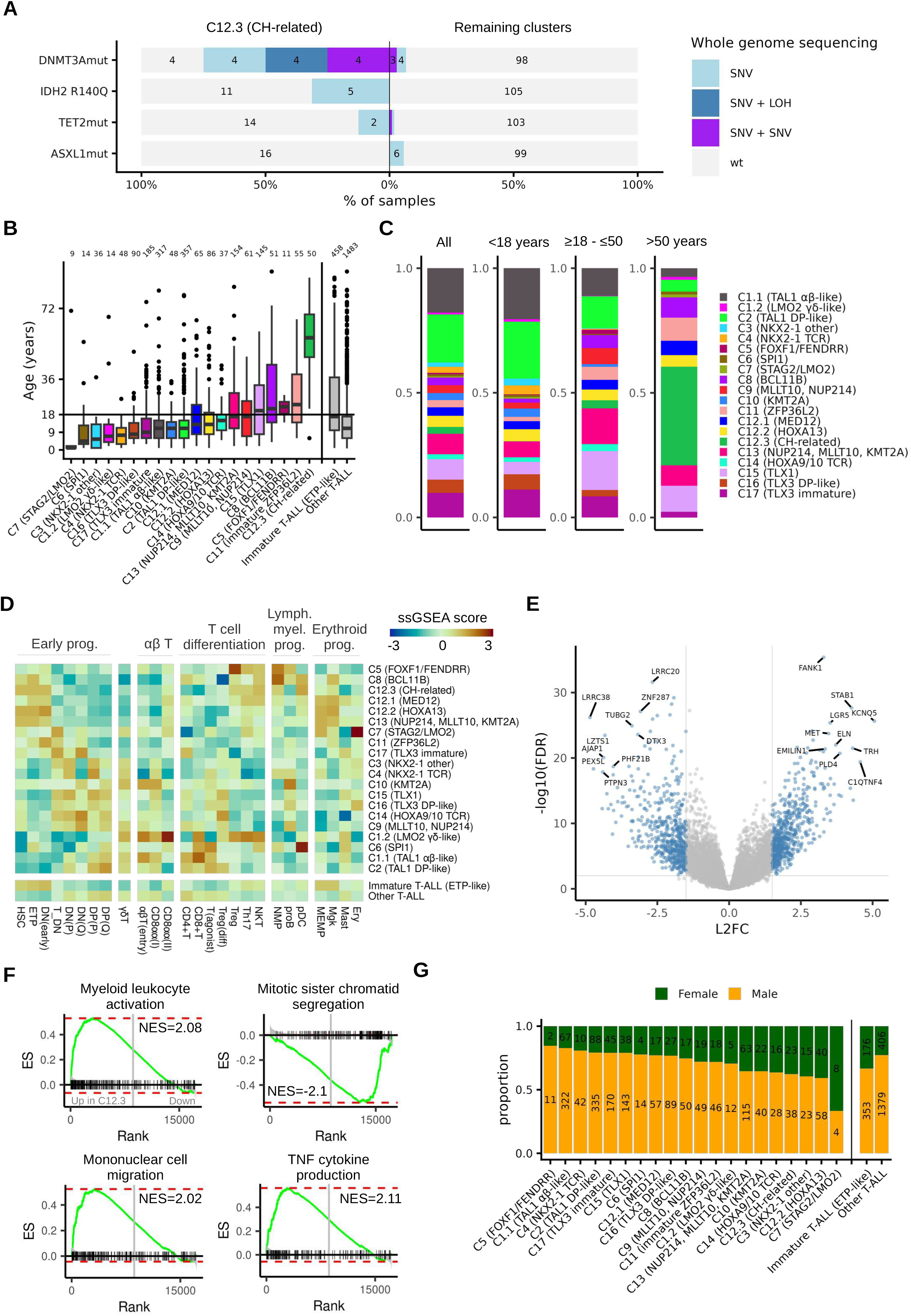
A novel clonal hematopoiesis related T-ALL subtype occurs frequently in patients older than 50 years and resembles immature T cell developmental stages. (**A**) WGS data from an unselected adult cohort was used to determine the frequency of mutations in genes frequently involved in clonal hematopoiesis in samples with cluster C12.3 assignment vs. samples of the remaining cohort. Color codes indicate the presence of a single nucleotide variant (SNV) alone or in combination with loss of heterozygosity (LOH) or the presence of two SNVs in the same gene. (**B**) Age distribution across subtypes. Numbers above the boxplots indicate patients with available age information per subtype. (**C**) Stacked bar charts with subtype frequency in all samples and across different age groups. (**D**) Heatmap with single sample gene set enrichment analysis (ssGSEA) using gene sets specific for T-cell developmental stages. The gene sets were generated from a human thymus single cell atlas. Results were averaged and z-normalized across T-ALL subtypes. Cell types were grouped into six broader groups (‘early T-cell progenitors’, ‘γδ T’, ‘αβ T’, ‘T differentiation’, ‘lymphoid and myeloid progenitors’ and ‘erythroid progenitors’). (**E**) Volcano plot showing differentially expressed genes between C12.3 (CH-related) samples and other T-ALLs. Top 20 differentially expressed genes are labeled. (**F**) Iceberg plots showing gene set enrichment analysis for four selected pathways. (**G**) Patient sex distribution across subtypes. Sex information was available for 1,328 samples and predicted for 932 using ALLCatchR2. Number of females and males per subtype are indicated.

*DNMT3A* and *IDH2 R140Q* mutations are frequently found in early/undifferentiated leukemias and 60/61 C12.3 (CH-related) samples were predicted to be immature T-ALL (ETP-like) consistent with an early developmental origin. Immunophenotypically, the C12.3 cluster was enriched for ETP ALLs (32.3% versus 11.2%, *P*<0.01). To investigate underlying developmental trajectories for all subtypes, we defined gene sets for thymic and hematopoietic cell types derived from single-cell RNA-Seq data^28^ (**Supplementary Figure 14A**). The 26 annotated cell types (**Supplementary Table 14**) spanned the full T-cell differentiation continuum, from hematopoietic stem cells (HSCs), ETP and double negative (DN) to mature αβ and γδ T cells (**Supplementary Figure 14B**) with high discriminative power (**Supplementary Figure 14C**). C12.3 (CH-related) had a high enrichment to HSC and grouped next to C8 (BCL11B) samples, which are often lineage-ambiguous (**Figure 5D**). C12.3 (CH-related) also showed high enrichment for plasmacytoid dendritic cell (pDC)-related genes, but the highest pDC association was observed in the C6 (SPI1) subtype, that has been indicated to originate from a T-cell precursor with DC potential. ^10^ Cluster C7 (STAG2/LMO2) localized adjacent to these early subtypes but showed stronger resemblance to thymic DN and erythroid progenitor profiles. C1.2 (LMO2 γδ-like) showed highest γδ T-cell enrichment, whereas C2 (TAL1 DP-like) was more similar to DP stages but showed lower αβ T-cell stage enrichment than C1.1 (TAL1 αβ-like).

The immature nature of C12.3 (CH-related) samples was characterized by high expression of stem cell markers (*CD34* and *cKIT*), and *LGR5* and *MET* - both involved in stemness - were the most upregulated genes (**Figure 5E**, **Supplementary Table 15**). Top enriched pathways were related to myeloid / monocytic inflammatory response (**Figure 5F**, **Supplementary Table 16**) suggesting multilineage potential in C12.3 or involvement of chronic inflammatory signaling.

Despite the overall male bias in T-ALL, female patients were more frequent in C12.3 (CH-related) samples (37.7% versus 24.5%, *P*<0.05). This was consistent with the relative sex convergence observed in other immature subtypes C12.2 (HOXA13) and C13 (**Figure 5G**). However, the immature T-ALL (ETP-like) enriched subtypes C11 (ZFP36L2) and C12.1 (MED12) did not show this tendency.

### Immature T-ALL predictions correlate with progenitor-like subpopulations

Immature T-ALL (ETP-like) predictions were highly specific to clusters C11 (64/64), C12 (239/244), and C13 (170/178), but were also detected at lower frequencies in other clusters, including C10 (KMT2A; 9/62), C17 (TLX3 immature; 8/215), C9 (MLLT10, NUP214; 5/68), C1.1 (TAL1 αβ-like; 5/389), C1.2 (LMO2 γδ-like; 2/17), and C8 (BCL11B; 1/67) (**Supplementary Table 1**).

Because bone marrow progenitor (BMP)-like subpopulations in T-ALL^29^ and pan-HSPC-like programs across leukemias have recently been linked to chemoresistance and poor outcomes,^30^ we performed enrichment analyses for the corresponding signatures. For 958 of 2,049 samples, AUCell enrichment scores from the original study were available and showed high concordance with our GSEA-derived BMP-like scores (ρ = 0.939 for the 119-gene signature and ρ = 0.927 for the 17-gene signature; **Figure 6A**). ALLCatchR2 immature T-ALL (ETP-like) prediction scores correlated with BMP-like/pan-HSPC enrichment and with ETP/DN(early) developmental signatures (**Figure 6A**; **Supplementary Figure 15A**). Samples predicted as immature T-ALL (ETP-like) consistently showed high BMP-like enrichment, suggesting agreement between machine-learning-based and gene set-based measures of early progenitor-like leukemia states (**Figure 6B**). C8 (BCL11B) samples represented a notable exception with high progenitor enrichment scores but no immature T-ALL (ETP-like) call, consistent with previous descriptions. Minimal overlap among gene sets for BMP-like, pan-HSPC, T-cell developmental, and immature (ETP-like) classification suggests that they capture complementary biological dimensions (**Supplementary Figure 15B**); therefore, BMP-like and pan-HSPC scores were added to the ALLCatchR2.

**Figure 6.**
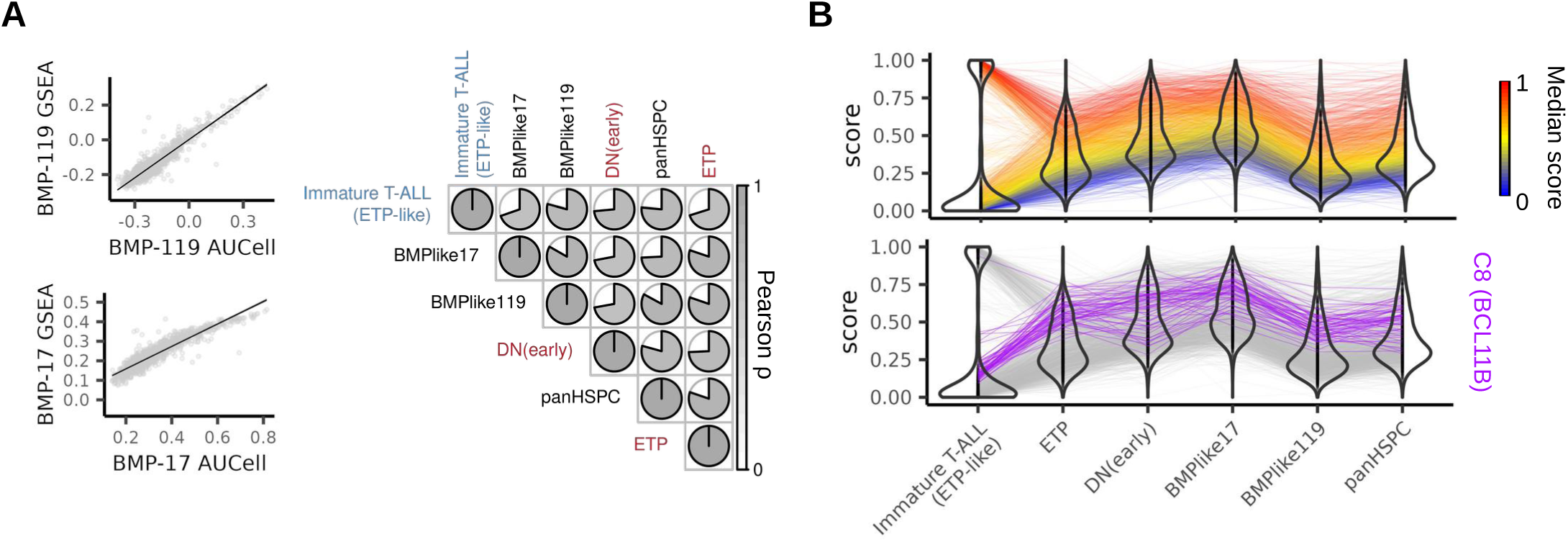
Immature T-ALL (ETP-like) enriched subtypes correlate with signatures from progenitor-like leukemia cells. (**A**) Pearson correlation analysis of recently published gene sets for bone marrow progenitor (BMP)-like leukemia subpopulation and hematopoietic stem and progenitor populations in acute leukemia (panHSPC). For 958 samples, the published BMP-like AUCell scores for gene sets including 119 and 17 genes could be compared to results from the ssGSEA analyses in this study. Enrichment scores for gene sets from published progenitor populations correlated with immature T-ALL (ETP-like) machine learning predictions (blue) and enrichment scores for ETP and DN(early) gene sets extracted from thymus single cell reference (red). (**B**) Predicted immature T-ALL (ETP-like) samples have high scores in the enrichment analyses. Lines are colored according to the average value across the six scores. C8 (BCL11B) samples are not part of the immature T-ALL (ETP-like) definition but show high enrichment scores for early T cell developmental stages and progenitor subpopulations.

## Discussion

Heterogeneous molecular subtypes have been proposed for T-ALL^1–7^ with a recent comprehensive pediatric/young adult landmark analysis providing 15 subtype definitions linked to developmental origins and clinical outcomes.^10^ This level of molecular granularity surpasses the resolution of conventional cytogenetics or immunophenotyping. Gene-expression profiles recapitulate the functional consequences of genomic drivers. In B-ALL, they have been implemented as independent diagnostic definitions (e.g. *BCR::ABL1*-like^31^ or PAX5alt^32^ B-ALL) and classifier tools such as our ALLCatchR achieve >0.95 accuracy in assigning individual samples to one of 22 subtypes^24,25^ consistent with transcript-inferred genomic drivers in 80% of cases.^22^ T-ALL poses specific challenges to this concept because (i) more than half of the drivers affect non-coding genomic regions difficult to identify in transcriptomic profiles^10^ and (ii) because consensus definitions of gene-expression subtypes are lacking and (iii) gene-expression profiles in T-ALL are less distinct. Consequently, the integrated analysis of genomic and transcriptomic data for the identification of genomic alterations and disease subtypes in diagnostics has been proposed^33^. However, available approaches for predicting T-ALL gene-expression subtypes remain limited.

Here, we leveraged the resolution of 2,314 aggregated T-ALL transcriptomes from 15 cohorts to propose a cohort- and age-spanning consensus of gene-expression clusters, based on an extended unsupervised analysis of high-blast cases. Supervised analyses were used only to define rare but clinically important LMO2 γδ-like T-ALL and an overarching ‘immature ETP-like’ definition. Comparison with subtypes assigned in the original publications indicated largely overlapping definitions despite partly heterogenous nomenclature. Clusters C6 (SPI1), C8 (BCL11B), C9 (MLLT10, NUP214), C10 (KMT2A), and C15 (TLX1) had been described in several cohorts across different age groups and previously described subdivisions^10^ within *NKX2-1* (C3, C4) and *TLX3* (C16, C17) were recognized as independent clusters in our analysis. In contrast, C1.1 (TAL1 αβ-like), C1.2 (LMO2 γδ-like), C2 (TAL1 DP-like) and C7 (STAG2/LMO2) had been described solely in the large pediatric landmark analysis.^10^ Mapping of genomic drivers available from 11 cohorts confirmed a cross-cohort consistency between drivers and gene-expression subtypes in T-ALL. Nevertheless, driver enrichments were not unambiguous, indicating that gene expression reflected here also the developmental origin. Although newer developments aim to integrate of whole genome sequencing and gene-expression profiling in T-ALL,^34^ we demonstrate the feasibility of cross-cohort gene-expression based subtype definitions. Importantly, gene-expression clusters in T-ALL partly overlap, especially in closely related clusters such as C1.1 (TAL1 αβ-like) and C2 (TAL1 DP-like) or C16 (TLX3 DP-like) and C17 (TLX3 immature). Moreover, our subtype definitions should not be considered static. We observed that the same driver genes can occur across multiple clusters, as well as indications of further subgrouping within clusters. To account for uncertainty, we trained our machine-learning predictor to perform quantitative multi-class predictions that help clarify the robustness of subtype assignment in a sample that may lack an identifiable genomic driver. Integration of more samples will likely enable a higher-resolution refinement of subtype definitions. One limitation of gene expression–based subtyping in general is its limited ability to distinguish between driver alterations that produce similar downstream transcriptional effects or to identify drivers with only a reduced impact on gene expression, even when such distinctions may be therapeutically relevant. Relatedly, while our quantitative multi-class framework flags uncertain cases by reporting candidate subtype calls, drivers with a stronger transcriptional footprint will inevitably dominate the final classification and may obscure co-occurring drivers of potential therapeutic relevance. Furthermore, gene expression–based resolution of rare genomic subtypes such as NKX2-5, or NUP98 remains constrained by their scarce representation in the reference cohort, and continued expansion of cross-cohort data sets will be required to fully capture these and other emerging entities. These considerations highlight that ALLCatchR2 is best applied as a complement to - rather than a substitute for - integrated genomic and transcriptomic analyses, particularly when the goal is identification of actionable driver alterations in a clinical context. Although our cohort spans the full age range, adults ≥40 years remain underrepresented (9.6% of samples with age information), and we cannot exclude that subtype specificities of older-adult T-ALL may still be missed. This is particularly relevant because some leukemic drivers are selected in an age-specific manner. However, in the unsupervised analysis, no subclusters were driven entirely by age independently of other defining features such as drivers. Together with the continuous AYA-to-adult gradient in subtype composition, this supports the cross-age validity of the proposed cluster definitions.

Importantly, we identified a novel gene-expression subtype enriched for the CH-associated mutations *DNMT3A* and *IDH2 R140Q*. This cluster was nearly exclusively observed in adults, accounting for 39.5% of T-ALL in older patients. A CH origin of T-ALL cases involving *DNMT3A* or *IDH2* mutations^35^ and the overall presence of *IDH1* and *IDH2* mutations in T-ALL^36^ had previously been described. Our analysis, however, demonstrates a distinct cluster of shared gene regulation that defines a specific CH-related T-ALL subtype. In line with published evidence^35,36^ and the biologically expected stem-cell near origin, CH-related T-ALL showed proximity to stem-cell and early T-cell developmental stages. This places CH-related T-ALL close to MPAL or even AML. Immunophenotyping warrants special attention here, because high expression of stem cell markers and frequent myeloid co-expression represent challenges for a clear lineage classification. Anecdotal reports have successfully applied AML-based hypomethylating agent/venetoclax combinations or enasidinib in *IDH2*-mutated T-ALL^37,38^ while *IDH1/2* mutated T-ALL cases had dismal outcomes on standard therapy underscoring the prognostic and potentially also predictive role of our novel subtype definition.

To facilitate systematic subtype assignment of T-ALL gene-expression profiles, we implemented our newly established models into ALLCatchR2. Building on the established B-ALL framework^24^ and by jointly modeling lineage, blast fraction, and subtype, ALLCatchR2 allows subtype prediction of ALL RNA-Seq samples independent of prior lineage assumptions. This provides a novel foundation for systematically harmonized T-ALL annotation across studies and paves the way for incorporation of molecular subtypes and developmental states into future research and also clinical stratification. Together with machine-learning models developed for B-ALL, ALLCatchR2 allows the identification of 42 age-spanning ALL subtypes, including the novel CH-related T-ALL subtype, enabling systematic analysis of ALL molecular subtypes and their developmental underpinnings. ALLCatchR2 is freely available as an R-package through https://github.com/ThomasBeder/ALLCatchR2.

## Data availability

Newly generated data sets are available from the European Genome Phenome Archive (EGA) under the accession number EGAS50000001829. Gene-expression count data form our own cohort^7^ is available Zenodo (10.5281/zenodo.20351819). Public available gene expression data was obtained from different sources. Pölönen et al.^10^ data are available from the Gabriella Miller Kids First Data Resource Center (Kids First DRC) for the Kids First: T-cell Acute Lymphoblastic Leukemia (KF-TALL) study. Data sets available in the NCBI Gene Expression Omnibus (GEO) repository under accession numbers GSE110633, GSE110636, GSE137768, GSE207057, GSE216117, GSE124824, GSE228632 and GSE78785 were accessed at https://www.ncbi.nlm.nih.gov/geo/. Bio-Med Big Data Center (BMDC) data sets were obtained from https://www.biosino.org/node/browse?keyword=OEZ00010057 and https://www.biosino.org/node/browse?keyword= OEZ00014089.

## Acknowledgements

This study was funded by the Deutsche Forschungsgemeinschaft (DFG; German Research Foundation) project number 444949889 (KFO 5010 Clinical Research Unit “CATCH ALL” to N.W., S.B., A.M.H., M.B., C.D.B., M.N. and L.B.) and project number 413490537 (Clinician Scientist Program in Evolutionary Medicine to D.B.), Deutsche José Carreras Leukämie-Stiftung (DJCLS 08R/2024 to L.B. and M.B.) and Deutsche Krebshilfe (70115443 to L.B.). L.B. was funded by the Faculty of Medicine of Kiel University within the Advanced Clinician Scientist Program. The GMALL trial 07/03 and the GMALL trial 08/13 were funded by Deutsche Krebshilfe. This work used data generated by the TARGET (Therapeutically Applicable Research to Generate Effective Treatments) initiative, supported by the National Cancer Institute. The large language model ChatGPT (OpenAI, USA) was used for language editing without interfering with any content of the manuscript.

