## Supplementary Methods for "The machine-learning classifier ALLCatchR2 identifies 20 T-ALL subtypes across cohorts and age groups"

### Supplementary Material and Methods

#### RNAseq

RNAseq was performed in 77 samples as described previously[1]. In short, RNA was extracted form bone marrow / peripheral blood mononuclear cells after Ficoll densitiy gradiet centrifugation and used for polyA-enrichment based library preparation (TruSeq RNA Library Prep kit (Illumina©, San Diego)) for stranded mRNA. Libraries were sequenced on Illumina NovaSeq platforms with 2 × 100-paired-end reads aiming for 30 million reads per sample. RNA-seq data analysis was performed using IntegrateALL[2] including read alignment to GRCh38.83 with STAR, Fusion calling (ARRIBA[3] and FusionCatcher[4]) and raw single-nucleotide variant calling (GATK, Genome Analysis Toolkit).

#### T-ALL blast count classifier

T-ALL sample blast counts were available for 3 cohorts (n=1008,[5] n=191,[1] and n=118[6]). The classifier was established on the 1008 samples from Pölönen et al. using an ensemble of linear regression (enet and glmnet) and random forest models (ranger and rf) implemented in caret.[74]Training was performed in an inner 5-fold cross-validation and initially based on the 3000 most variable expressed genes. To define the most important genes for each classifier we used the varImp function[7] and identify_outliers from the R package rstatix. Next the classifiers were trained using the most important genes for each machine-learning method and blast count predictions averaged for ensemble classification. Testing of the classifier was performed using two independent hold-out cohorts with n=191[1] and n=118[6] samples. Normalization of the original counts was performed by $\log_2$(count + 1) transformation, followed by z-score scaling for each sample individually to avoid data leakage.

#### Acute lymphoblastic leukemia lineage classifier

In order to assign samples with unknown lineage-information, we trained a lineage classifier based on two cohorts that contained both B-ALL and T-ALL data. By using these data set we avoided combination of technical/processing effects and lineage signals entering the classifier training. GMALL data comprised 774 B-ALL and 307 T-ALL samples from published resources[1,2,8] but also 77 new T-ALL samples. The second data set from Yu et al.[9] comprised 110 B-ALL and 32 T-ALL samples. Classifier training, was performed using caret[7] package and an ensemble classifier established using the algorithms svmLinear3, svmLinearWeights2, ranger, rf and LogitBoost. Training was performed in an inner 10-fold cross-validation scheme for hyperparameter optimization. Least Absolute Shrinkage and Selection Operator (LASSO)[10] and α parameter 0.1 was applied for feature selection using the glm-net[11] package.

### Unsupervised T-ALL gene expression subtype identification

In total, 2,049 known and predicted T-ALL samples were included in the analysis. Expression data were harmonized across cohorts based on 17,361 genes detected in all datasets, and batch correction was performed using ComBat-seq[12] as implemented in the sva[13] package to mitigate technical differences between cohorts. Low-variance genes (n = 630) were then removed using the nearZeroVar function in caret[7] with freqCut = 97/3. Counts were normalized using $\log_2$(count + 1) transformation, followed by z-score scaling across samples. Although ComBat-seq effectively reduces technical variance while preserving biological signal, unsupervised methods remain sensitive to residual batch effects that can still influence clustering. In addition, sex-linked gene expression introduced unwanted structure. Therefore, genes associated with sex (n = 349) and cohort (n = 870) were removed based on absolute Spearman correlation thresholds of 0.1 and 0.25, respectively. After filtering, 15,512 genes remained for unsupervised analysis.

The unsupervised analysis was performed on 1,724 of the 2,049 samples that had predicted blast counts >70%. In addition one known tumor-microenvironment–enriched sample[5] with a predicted blast count of 70.3% was excluded. These preprocessing steps rigorously minimized batch effects and revealed clear gene expression clusters (**Figure 1A**). Unsupervised grouping of samples was performed using the 400 most variably expressed genes. Sample-to-sample distances were derived from 24 UMAP analyses,[14] representing all combinations of the parameters *n_neighbors* = 3, 5, 10, 20 and *min_dist* = 0.0125, 0.05, 0.1, 0.2, 0.5, and 0.8. For each UMAP run, pairwise distances between samples were computed, z-scaled, and then averaged across all 24 parameter settings to obtain a consensus distance matrix. This final distance matrix was subjected to hierarchical clustering using the Ward.D2 method implemented in *hclust*. The resulting dendrogram was progressively partitioned using *cutree* at each junction from 1 to 39 clusters. At each splitting level, robustness of the clusters was determined by the ability of a linear SVM classifier to reproduce the corresponding cluster. If Cohen's κ dropped below 0.8, the cluster was considered unstable, and we reverted to the previous clustering level before the split as the last stable cluster separation. This analysis revealed 17 main T-ALL gene expression clusters and final robustness was determined with an ensemble classifier (svmLinear3, svmLinearWeights2, ranger, rf and LogitBoost) and 10-fold randomized stratified cross-validation. Training was performed in an inner 10-fold cross-validation scheme and LASSO with α = 0.3 for feature selection. During training, class imbalances were adjusted using synthetic minority over-sampling technique (SMOTE).[15] The same approach was applied to all subsequent machine-learning classifiers unless stated otherwise.

**Subtype-specific refinement and cluster overarching definition of immature T-ALL**

The initial unsupervised analysis did not fully capture the diversity of T-ALL expression profiles. Given its clinical relevance, the recently described ETP-like subtype[5] was incorporated into the classification framework. Samples from our aggregated dataset were clustered using same UMAP parameter-combination strategy as described above but the top 1,000 differentially expressed genes between known ETP-like (n=132) and non–ETP-like (n=665) cases. This resulted in a highly distinct cluster enriched for ETP-like (**Supplementary Figure 7A**) and cluster robustness was determined by machine-learning.

Clusters C1 (enriched for TAL1 αβ-like and LMO2 γδ-like) and C12 (enriched for *MED12*, *HOXA13*, *ZFP36L2*, *DNMT3A*, and *IDH2* alterations) showed additional internal heterogeneity and were therefore analyzed separately. To distinguish the LMO2 γδ-like and TAL1 αβ-like subgroups within C1, differential gene expression analysis was performed, and 560 differentially expressed genes were used for unsupervised clustering. This analysis was conducted on n = 314 C1 samples using the same UMAP parameter-combination strategy as described above. Similarly, the 151 samples assigned to C12 were reanalyzed using the 400 most variably expressed genes and the same UMAP ensemble workflow. C1 and C12 subcluster robustness was determined by machine-learning.

**ALLCatchR T-ALL classifier training**

To implement the 20 T-ALL gene expression subtypes and the immature T-ALL (ETP-like) category in ALLCatchR, we used a core set of 1,567 representative samples from 14 cohorts. Core representative samples had predicted blast counts > 70% and were correctly assigned to one of the 17 main gene expression clusters in the initial analysis. The data were split into 1,364 training samples from 11 cohorts and 203 validation samples from three left-out cohorts. The composition of the training and validation datasets is summarized in **Supplementary Table 2**. For model training, original count data without batch correction was used and normalization of counts was performed for each sample separately by $\log_2$(count + 1) transformation, followed by z-score scaling to avoid data leakage. Then training was performed for each subtype with the ensemble classifier approach described above and an inner 10-fold CV. To account for class imbalance, we applied two rules: (1) for subtypes with fewer than 200 samples, the comparison class was sampled to 200 cases; and (2) for subtypes with more than 200 samples, both the subtype and the comparison class were down-sampled to 200 cases.

Cluster-specific prediction score cutoffs (**Supplementary Table 9**) were determined using the training set and were assigned to three confidence tiers: high-confidence, candidate, and unclassified. High-confidence predictions required a score exceeding the threshold at which samples from a given cluster were optimally separated from all other samples. This threshold was defined by the maximum Cohen's κ; if a range of scores yielded the same maximal κ, the mean of that range was selected. Candidate predictions were defined as scores ≥ 66% of the high-confidence cutoff. Samples that did not reach the candidate threshold for any subtype were designated as unclassified.

**Validation of CH-related SNVs of C12.3 predicted samples**

Genomic DNA from 10/14 GMALL samples predicted as *C12.3 (CH-related)* was analyzed using the Myeloid-NDC assay (Univ8 Genomics, Belfast, UK) according to the manufacturer's instructions. The assay enables integrated detection of sequence variants and structural variants associated with myeloid malignancies.

Sequencing was performed on an Illumina NextSeq 1000 platform using 2 × 100bp reads. Bioinformatic analysis was performed according to the manufacturer's recommendations using the Myeloid-NDC Analysis application on Illumina's BaseSpace Sequence Hub.

**Single cell reference of human T lymphopoiesis**

Single-cell RNA-seq data representing human T-cell developmental stages were obtained from a published reference[16] comprising two datasets. The first included 30,694 cells spanning early thymic development, and the second contained 76,994 cells from human thymic tissue. Cells were collected from a total of 27 pre- and postnatal donors and distributed across 112 samples generated through sorting into the 45P, 45N, CD3P, CD3N, CD137, 45NM, and EPCAM fractions. The resulting atlas encompassed 26 author-defined cell types (**Supplementary Table 14**).

Cells from both datasets were combined and processed using Seurat (v5.3.0)[17] with default parameters unless otherwise specified. After merging the count matrices (JoinLayers), data were normalized (NormalizeData), variable features were identified (FindVariableFeatures), and expression values were scaled and centered (ScaleData).

Dimensionality reduction was performed using PCA (RunPCA), followed by construction of the nearest-neighbor graph (FindNeighbors). Batch integration was conducted using Harmony[18] within Seurat (RunHarmony), with group.by.vars set to donor identity and library assay.

Gene sets specific to T-cell developmental stages were derived using LASSO regression (alpha = 0.1) applied to pseudobulk expression profiles computed for each cell type and sample. Genes with positive or negative regression coefficients were grouped for each cell type. Single-sample GSEA (ssGSEA) was performed using singscore[19] with these T-cell development gene sets.

#### Differential gene expression and gene set enrichment analysis

Differential gene expression analysis was performed using limma[20] and genes were ranked by their log2 fold change. Gene set enrichment analysis was performed using the fgsea[21] function and the following parameter: minSize = 25, maxSize = 250, eps = 0.0, using Gene Ontology (GO) Biological Process (BP) gene sets from MsigDB (v2025.1).[22]

#### TALLSorts

TALLSorts[23] was applied by converting gene symbols to Ensembl gene IDs using the EnsDb.Hsapiens.v86 R package and transposing the uncorrected count data of 2,049 T-ALL samples. To test correct tool implementation, we used count data from Verboom et al.,[24] comprising 85 samples from two cohorts, and 237 TARGET T-ALL samples, which have been used to train TALLSorts. Therefore, subtype predictions should match the ground-truth defined by the authors. This comparison showed that our predictions for 308/322 samples (95.65%) matched the ground-truth subtype labels used for TALLSorts training, confirming that the tool was applied correctly.
