## Supplementary Figures for "The machine-learning classifier ALLCatchR2 identifies 20 T-ALL subtypes across cohorts and age groups"

### Supplementary Figure 1

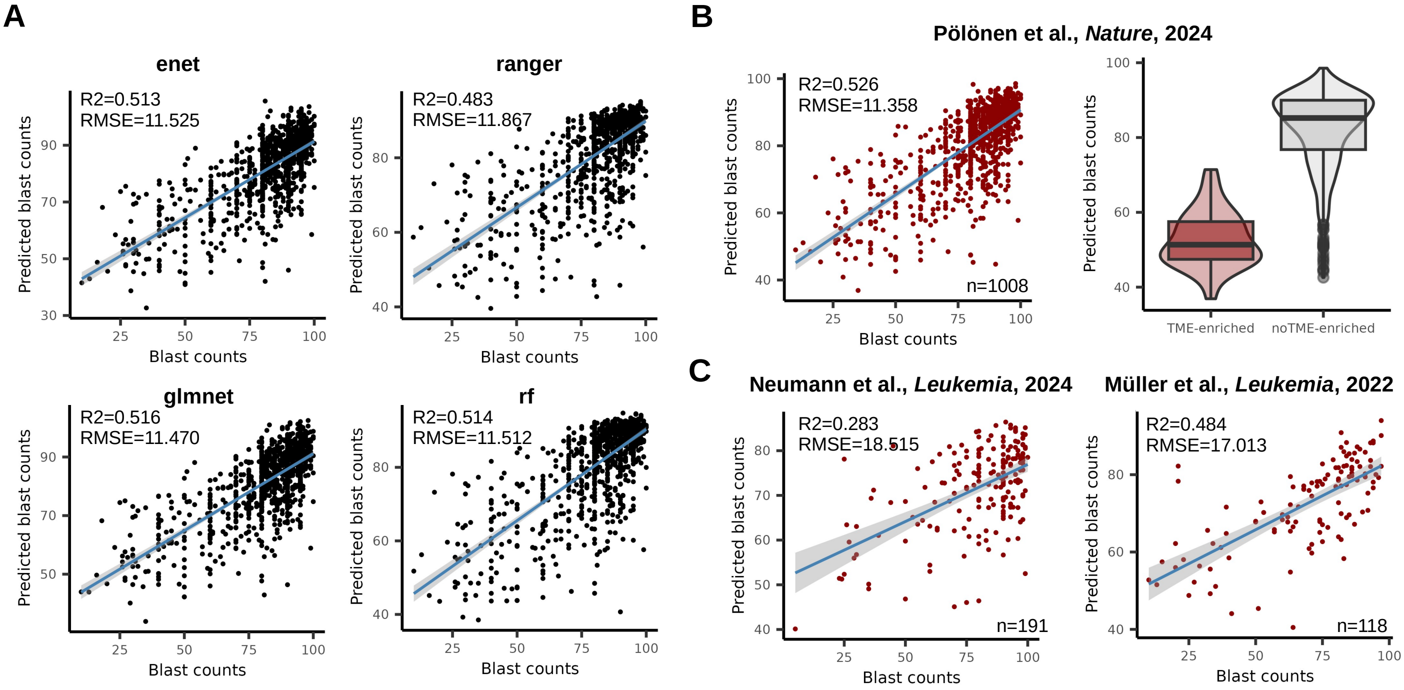


**Supplementary Figure 1. T-ALL blast count prediction using machine-learning regression models.** (**A**) Scatterplots showing blast counts and predictions for n=1,008 samples from Pölönen et al., Nature, 2024 that were used for model training. Two linear (enet and glmnet) and two random forest machine-learning models (ranger and rf) were used and all showed similar prediction performance as determined by Pearson R squared (R2) and root mean squared deviation (RMSE). Lines show linear regression and grey area the 95% confidence interval. (**B**) The final model is an ensemble of the models in (**A**) and samples belonging to the group tumor microenvironment (TME) enriched defined by Pölönen et al. had lower predicted blast counts than other T-ALL samples. (**C**) Blast count prediction for the testing data sets showed good prediction performance demonstrating model generalizability.

#### Supplementary Figure 2

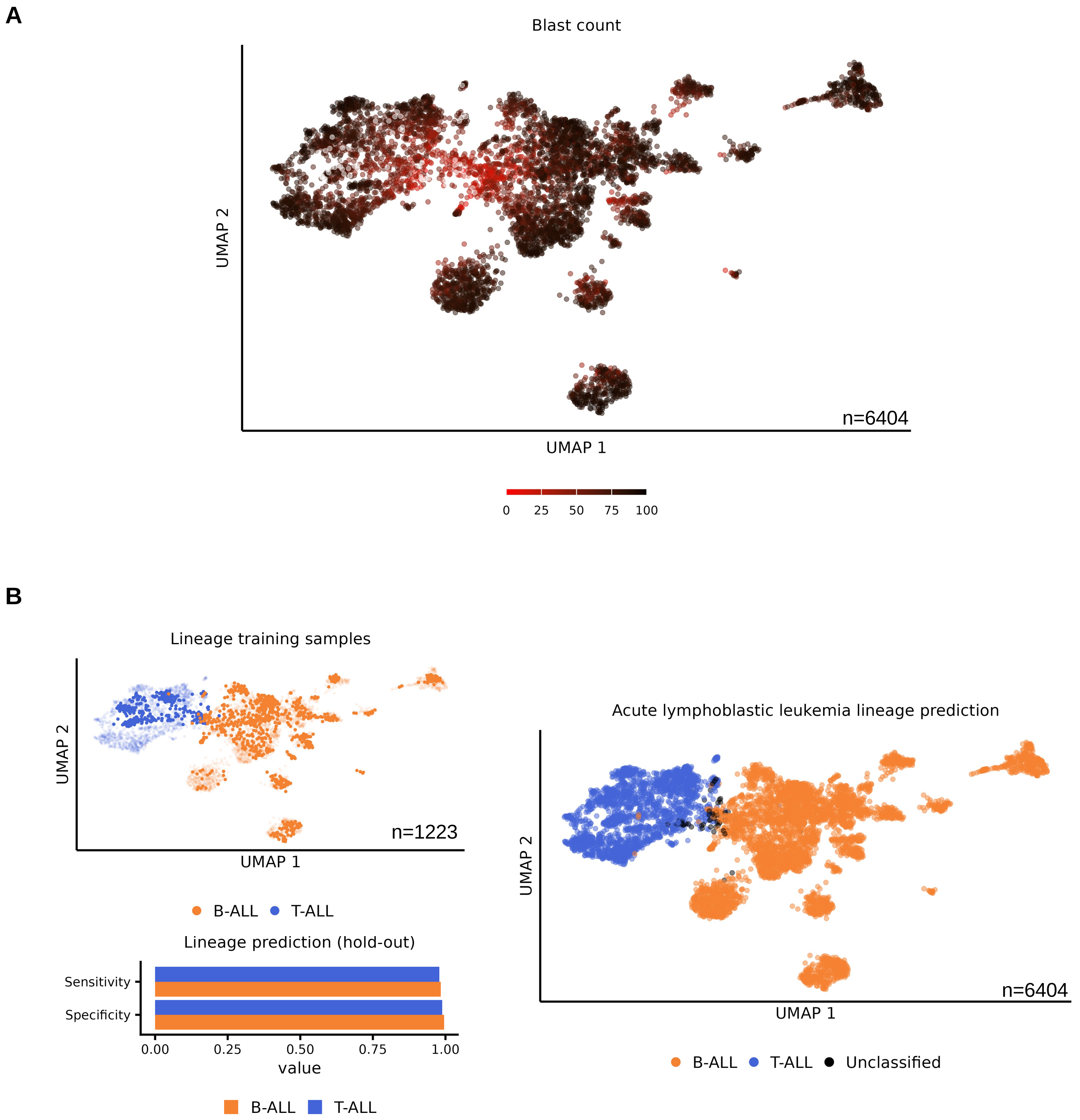


**Supplementary Figure 2. Acute leukemia lineage and blast count classifiers.** (**A**) UMAP analysis using the 2000 most variable expressed genes determined for 6,404 B-ALL and T-ALL samples. Samples are colored according to predicted blast counts and samples with low blast counts group at the center. (**B**) Same UMAP plot as in (**A**) but samples are colored according to disease lineage. Training samples are highlighted in the left plot and predictions shown in the right plot. Unclassified samples are enriched at the intersection between B-ALL and T-ALL and are associated with low blast counts. The barplot shows the prediction performance for the n=1667 T-ALL and n=3499 B-ALL hold-out samples not used for classifier training.

### Supplementary Figure 3

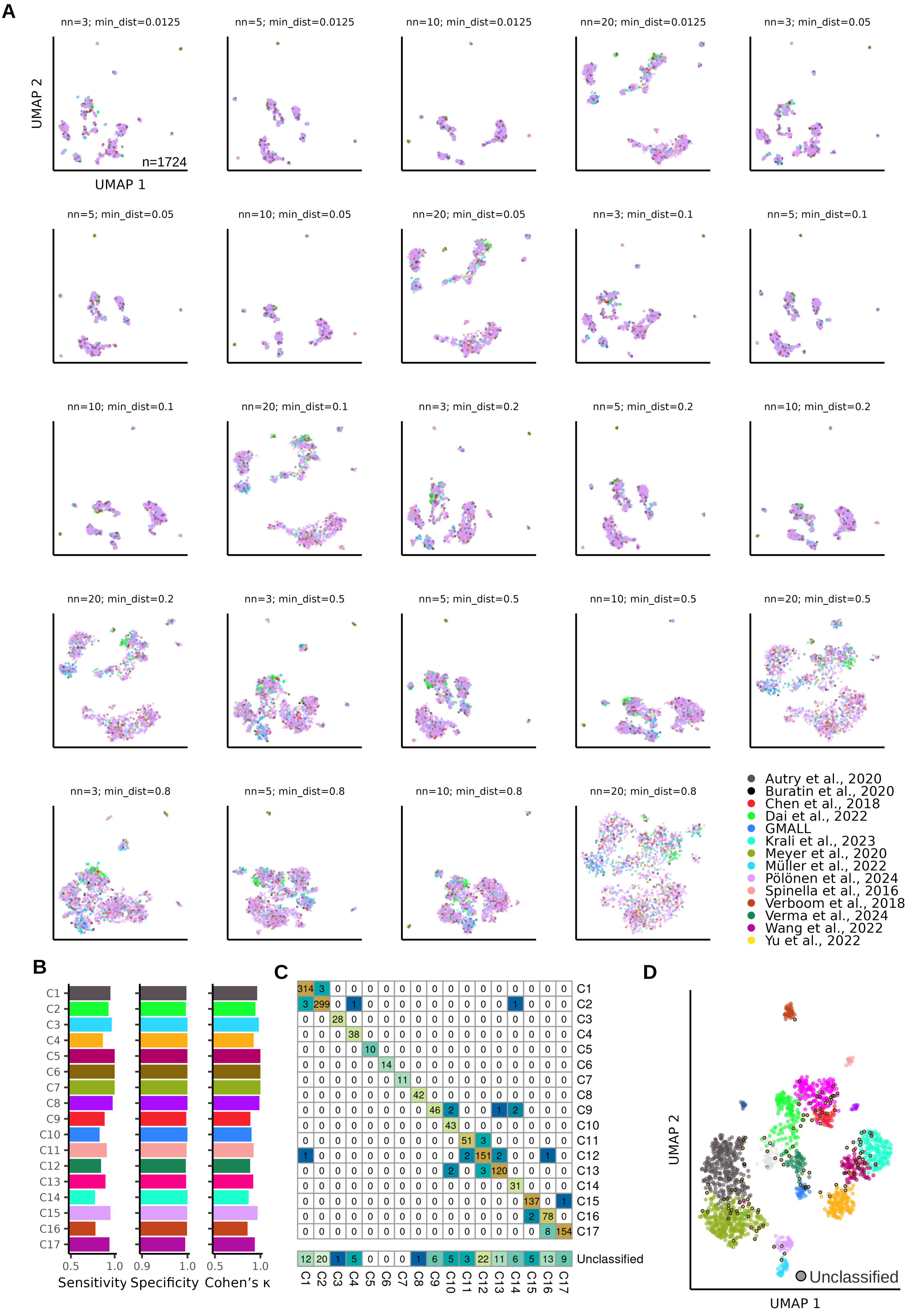

**Supplementary Figure 3. Unsupervised analysis of T-ALL samples revealed 17 gene expression cluster**. (**A**) UMAP plots of 1,724 samples with predicted blast counts >70% and systematic variations of minimum distance and number of neighbors. The analysis was based on the 400 most variable expressed genes and samples are colored according to contributing cohorts. (**B**) Prediction performance of main T-ALL gene expression clusters determined by sensitivity, specificity and Cohen's κ per cluster. (**C**) Confusion matrix showing number of samples in the 17 clusters in columns and corresponding machine-learning predictions in rows. The 1,567 core representative samples for each cluster were determined by the unsupervised analysis and correctly predicted (diagonal in heatmap). (**D**) UMAP plot based on 1,330 lasso genes shows predictions and distinct clustering of the 1,724 samples from 17 main gene expression cluster identified by the unsupervised analysis.

### Supplementary Figure 4

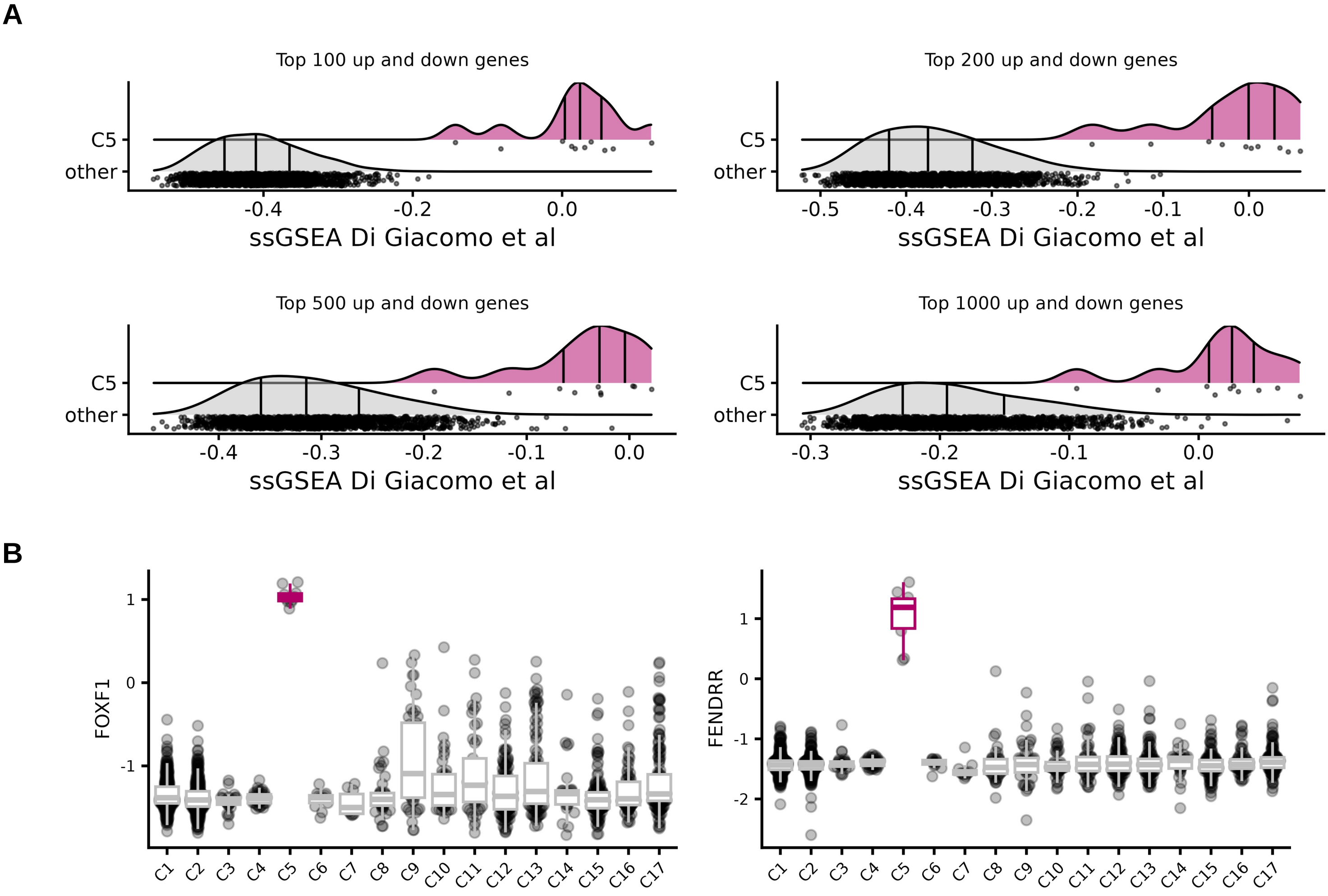


**Supplementary Figure 4. Gene expression cluster C5 corresponds to the recently defined FOXF1/FENDRR T-ALL subtype.** (**A**) Single sample gene set enrichment analysis using Di Giacomo et al. (Blood 2026) FOXF1/FENDRR differentially expressed genes. (**B**) Expression of marker genes for the new FOXF1/FENDRR across subtypes.

### Supplementary Figure 5

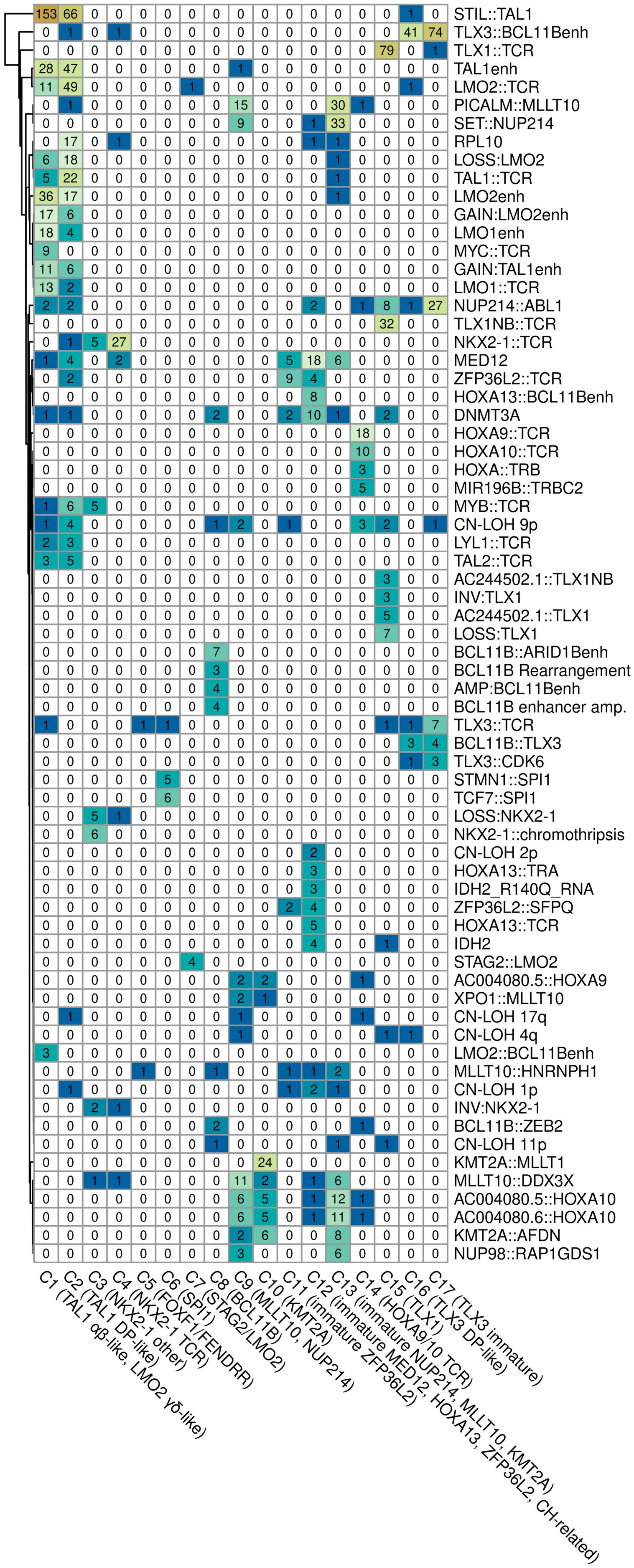


**Supplementary Figure 5. T-ALL gene expression clusters are characterized by presence of cluster specific drivers.** Heatmap showing drivers present in ≥3 samples across T-ALL main gene expression cluster.

### Supplementary Figure 6

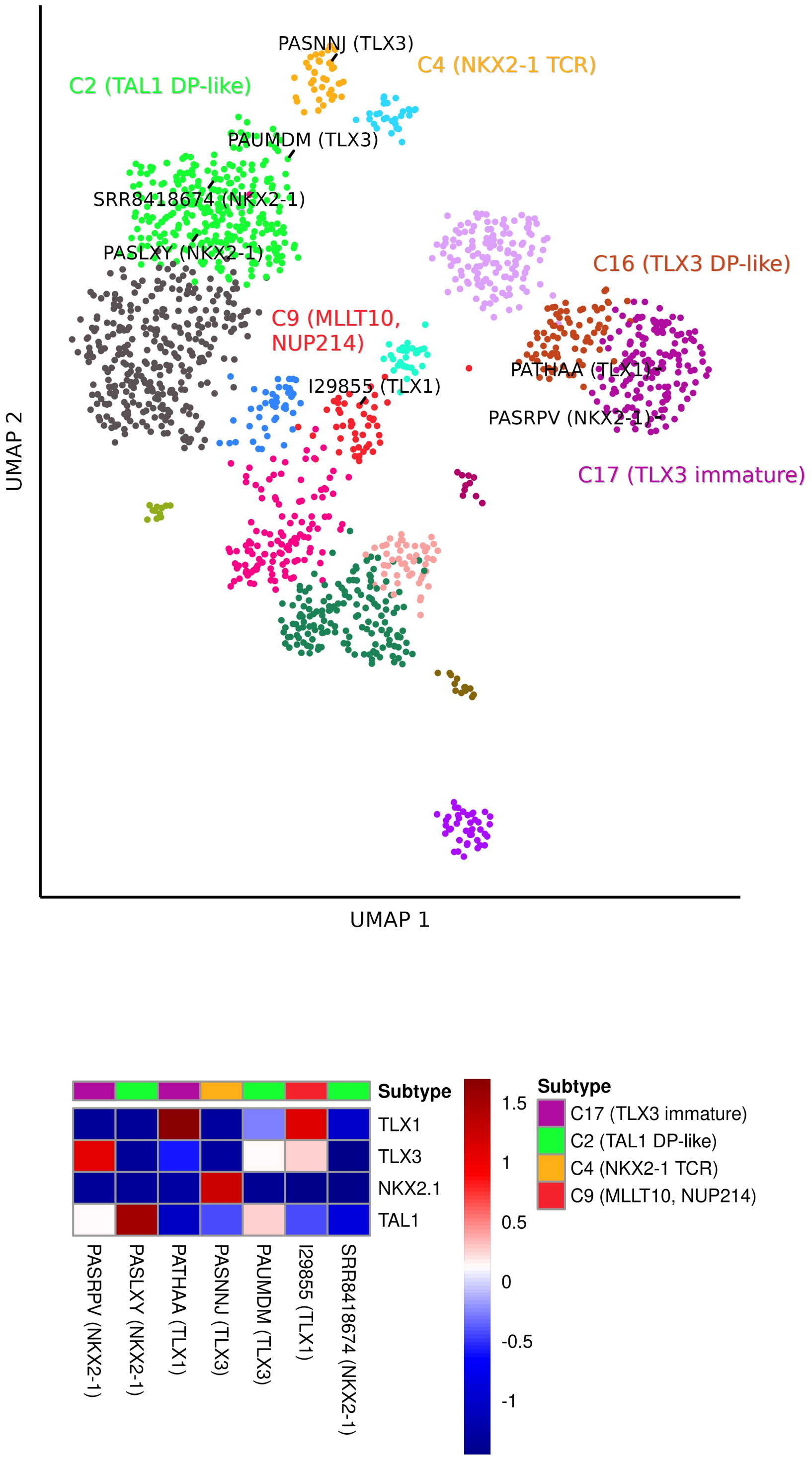


**Supplementary Figure 6. Discrepancies between published subtype assignments and gene expression subtypes. (A)** UMAP plot based on 1,330 lasso genes shows distinct clustering of 1,567 core representative samples of the 17 main gene expression cluster identified by the unsupervised analysis. Seven samples in the well defined NKX2-1, TLX1 and TLX3 subtypes in the original publications did not group in the corresponding gene expression cluster. Subtypes from the original publications are indicated in brackets next to the sample name. (**B**) Marker gene expression in these samples.

### Supplementary Figure 7

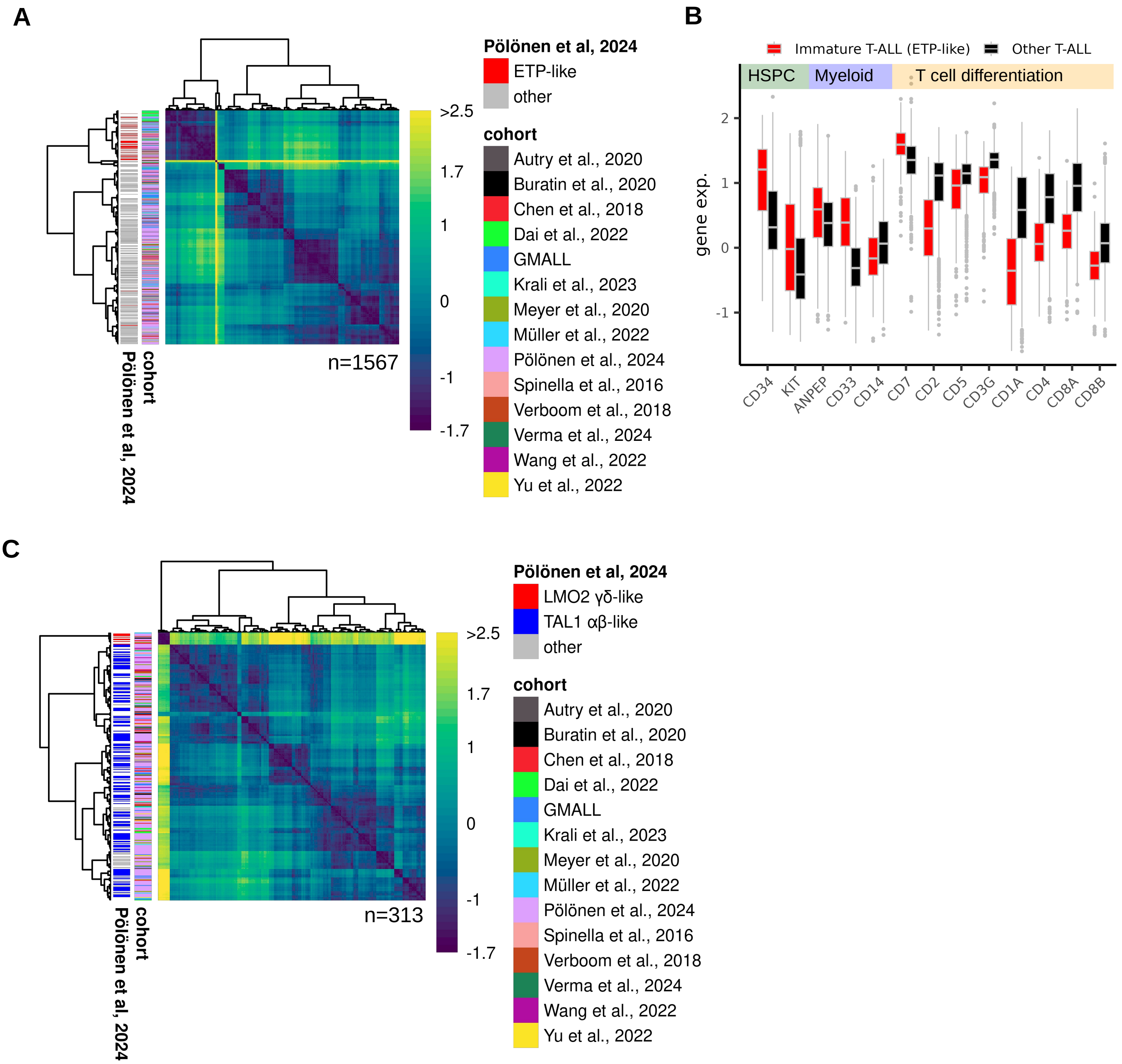


**Supplementary Figure 7. Identification of a cluster overarching definition of immature T-ALL and cluster 1 subgroups.** (**A**) Heatmap showing integrated sample-to-sample distances from 24 UMAP analyses of systematic variations of minimum distance and number of neighbors. Analysis was based on the 1000 most significant differentially expressed genes between defined ETP-like and non-ETP-like samples from Pölönen et al., Nature 2024. Hierarchical clustering revealed a distinct ETP-like enriched cluster independent of contributing cohorts. (**B**) Boxplots showing normalized expression of marker genes for hematopoietic stem and progenitor cells (HSPC), myeloid lineage and T cell differentiation in immature T-ALL (ETP-like) predicted samples compared to other T-ALLs. (**C**) Heatmap showing integrated sample-to-sample distances from 24 UMAP analyses for C1 (TAL1 αβ-like, LMO2 γδ-like) samples. Analysis was based on 560 differentially expressed genes between defined TAL1 αβ-like and LMO2 γδ-like samples from Pölönen et al., Nature 2024. Hierarchical clustering revealed two distinct groups independent of contributing cohorts.

### Supplementary Figure 8

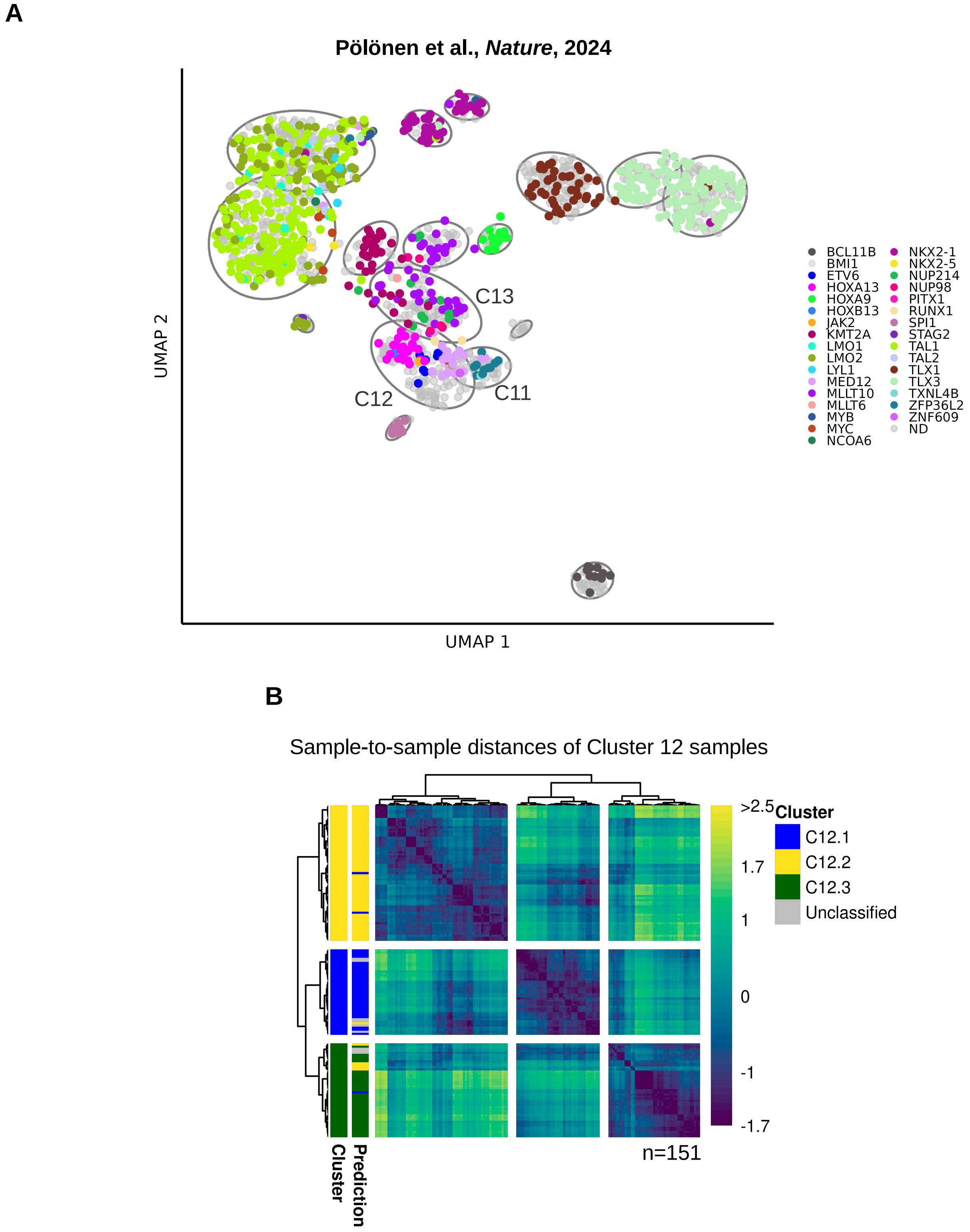


**Supplementary Figure 8. Gene expression subgroups within the cluster 12 were indicated by distinct driver enrichment**. (**A**) UMAP plot based on the 1,330 LASSO genes showing the 1,567 core representative samples. Samples are colored according to the driver gene annotations from Pölönen et al., Nature, 2024 on the left panel. Distribution of drivers in C12 indicated three distinct subclusters within C12 i.e., one characterized by *HOXA13*-rearrangements, one by *MED12* SNVs, and a third one without reported drivers in Pölönen et al. (**B**) Unsupervised analysis of C12 samples show three subgroups within C12 that could be predicted by machine learning.

### Supplementary Figure 9

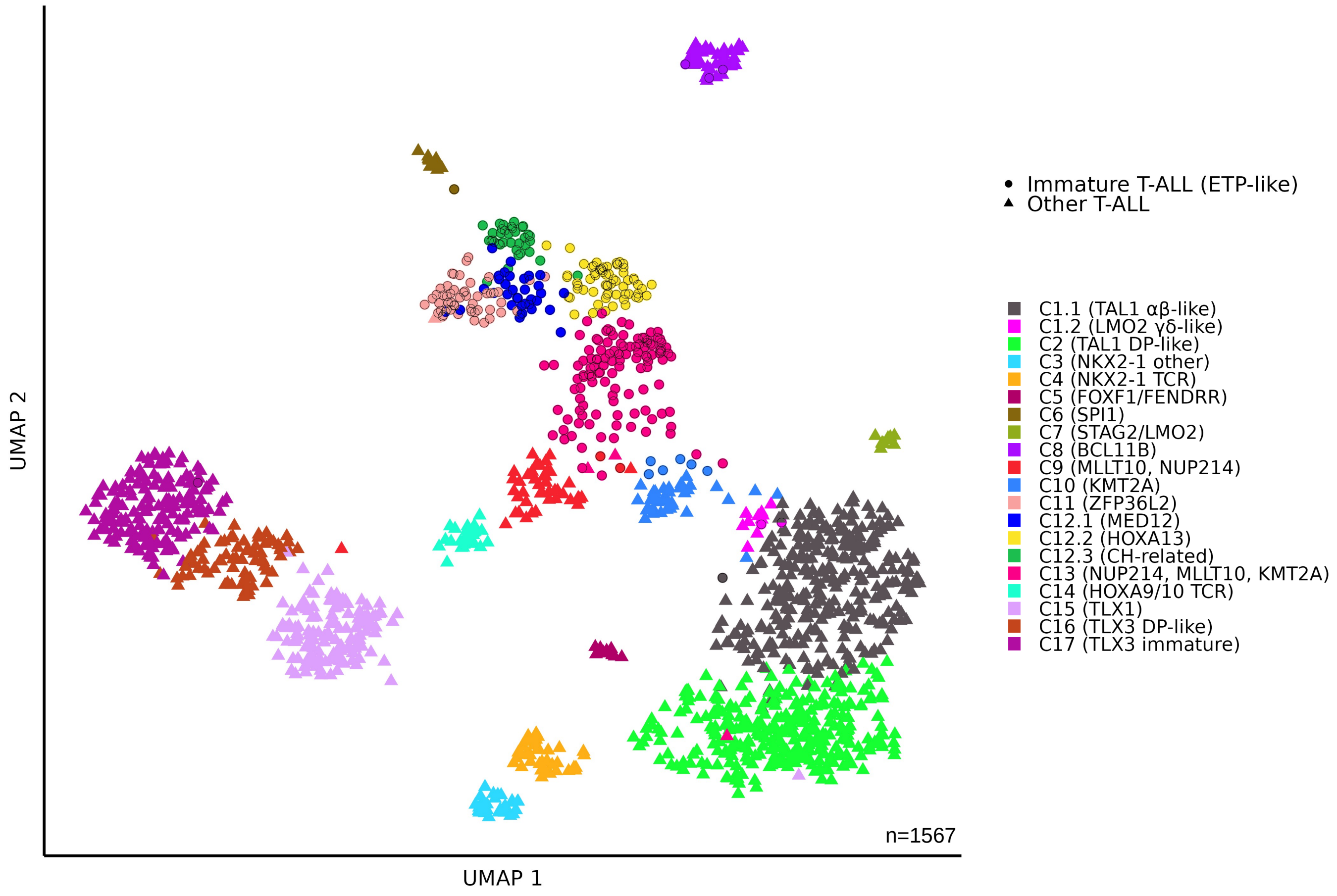


**Supplementary Figure 9. The 20 T-ALL gene expression subtypes and a subtype-overarching definition of immature T-ALL (ETP-like) are characterized by distinct gene expression profiles.** UMAP plot based on 1,689 LASSO genes selected for all 20 T-ALL gene expression clusters. Immature T-ALL (ETP-like) samples are indicated by icon shape.

### Supplementary Figure 10

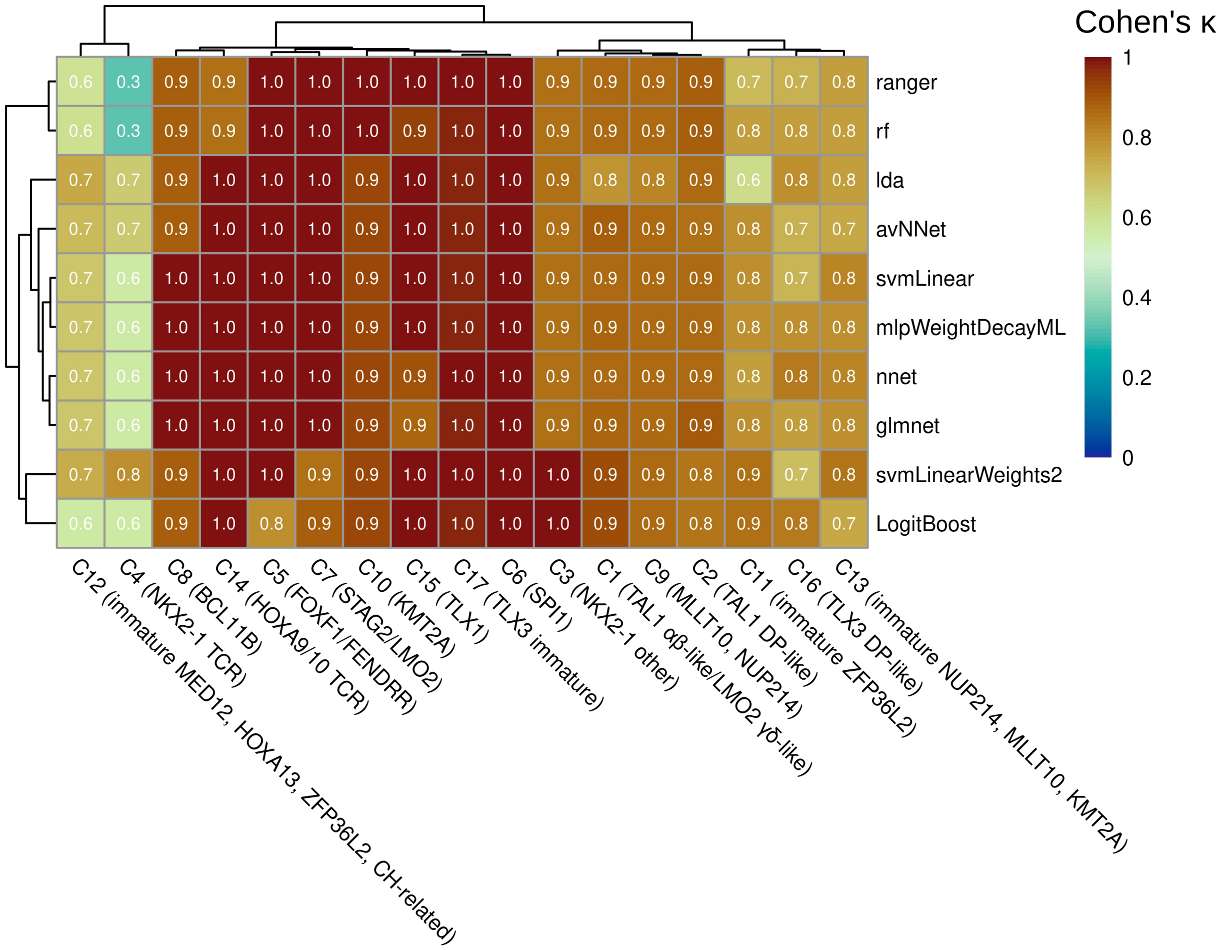


**Supplementary Figure 10. Prediction performance of different machine learning models for the 17 main T-ALL gene expression subtypes.** Heatmap showing Cohen's κ values of the 10 machine learning models for main gene expression clusters of the n=203 validation samples. Similar prediction performance was observed, but with certain subtype specific differences. The models for ALLCatchR2 T-ALL prediction were selected to account for this: rf and ranger had the highest prediction scores for C10 (KMT2A), svmLinear3 performed very well on C8 (BCL11B), svmLinearWeights2 performed best on C4 (NKX2-1 TCR) and both svmLinearWeights2 and LogitBoost performed best on C3 (NKX2-1 other). To calculate the performance for each subtype prediction scores > 0.5 were considered as prediction cut-off.

### Supplementary Figure 11

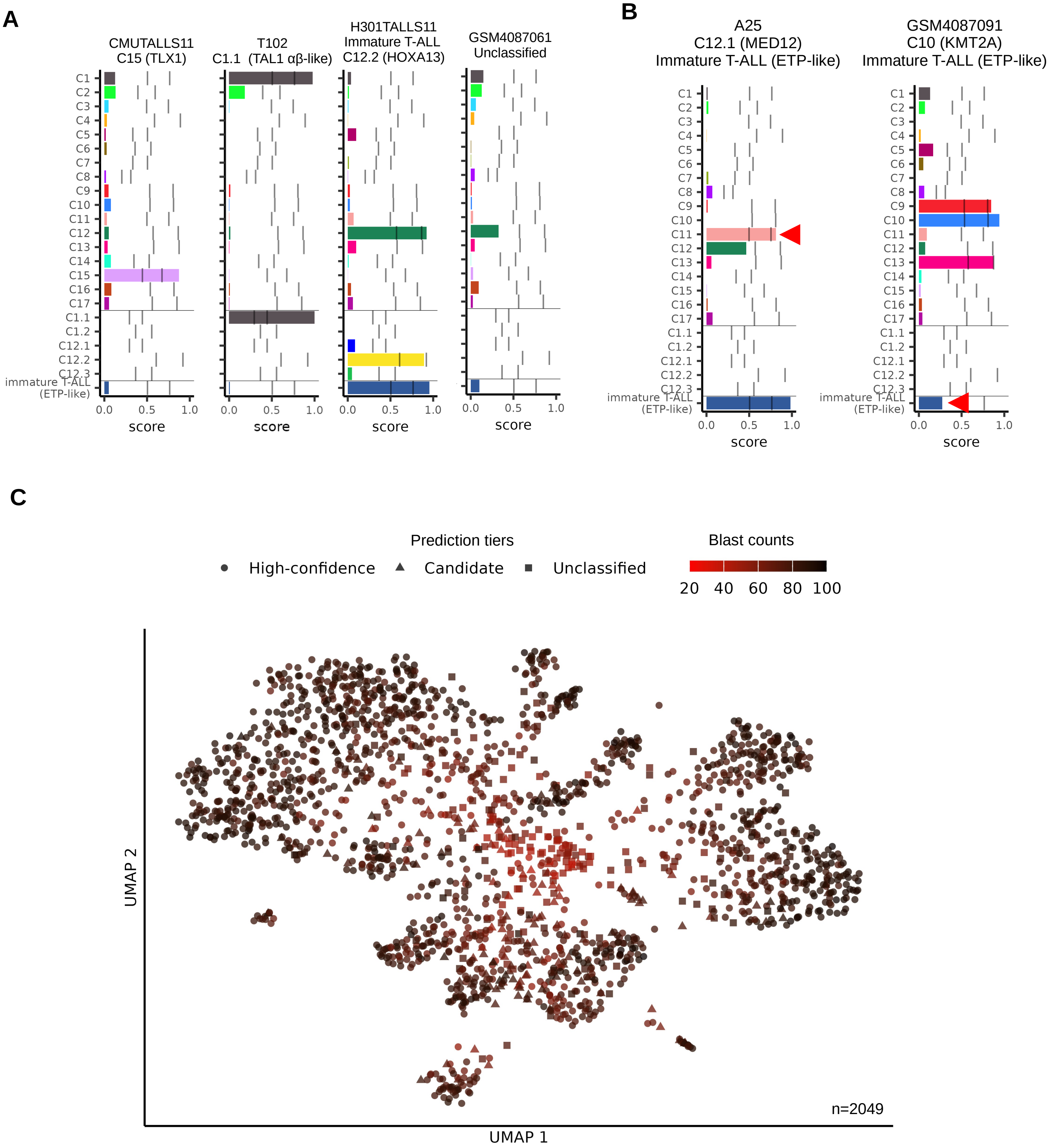


**Supplementary Figure 11. ALLCatchR2 for hierachical classification of T-ALL.** (**A**) Predictions scores for four samples illustrate hierarchical classification implemented in ALLCatchR2. If a sample did not reach candidate prediction cutoffs for any subtype it was considered unclassified. (**B**) Prediction scores for 2/203 misclassified samples from the validation set. Misclassifications are indicated by red arrow heads. Sample A25 belongs to cluster C12.1 (MED12) but was predicted as C11 (ZFP36L2) and sample GSM4087091 is an immature T-ALL sample that was misclassified as non-immature. (**C**) Same UMAP plot based on the 1,689 LASSO genes as in Figure 2C but colored according to the predicted blast counts per sample. Point shape illustrates the classification tier groups. Unclassified samples are enriched at the center of the plot and characterized by low blast counts.

### Supplementary Figure 12

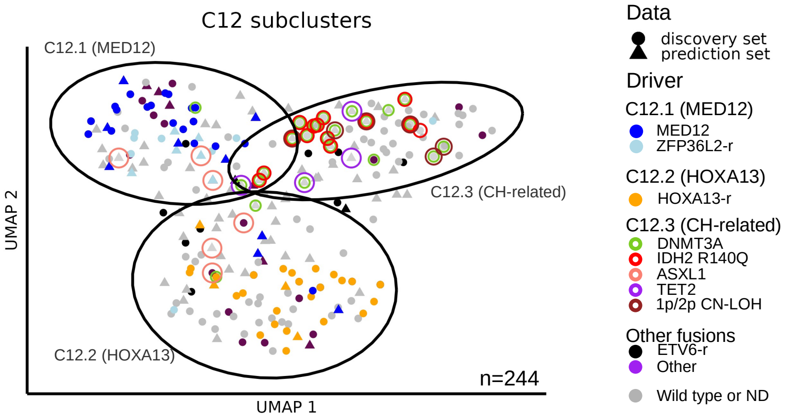


**Supplementary Figure 12. Samples predicted as C12.3 CH-related are enriched for *DNMT3A*, *IDH2* and *TET2* SNVs.** UMAP analysis using 322 LASSO genes and are enriched for SNVs in clonal hematopoiesis (CH) associated genes *DNMT3A*, *IDH2* and *TET2*. The ellipses in the plot are based on the 95% confidence region.

### Supplementary Figure 13

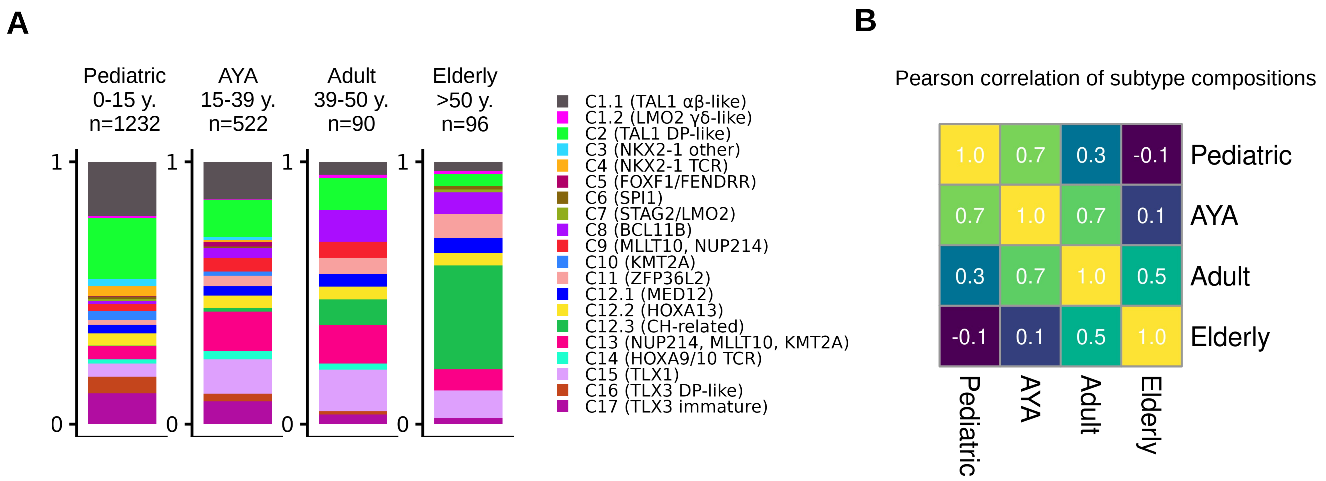


**Supplementary Figure 13. Distribution of T-ALL gene expression subtypes across age groups**. (**A**) The stacked bar plots on the left show the relative proportion of the subytpes within each age group. (**B**) The heatmap summarizes similarities in subtype composition between age groups using Pearson correlation coefficients. AYA, adolescent and young adult.

#### Supplementary Figure 14

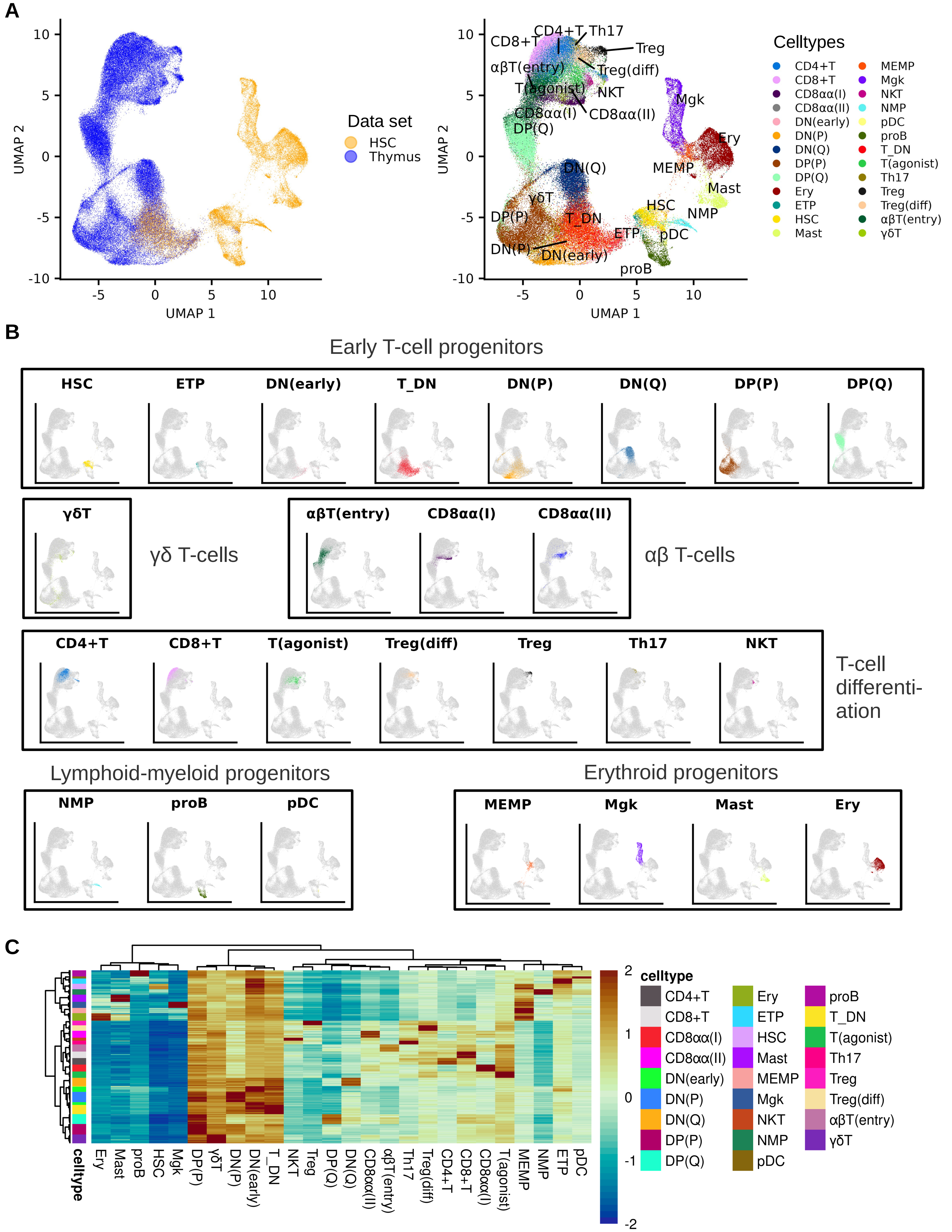


**Supplementary Figure 14. Gene set definition for T cell developmental stages using a single cell reference of human T lymphopoiesis.** (**A**) UMAP plot showing the two combined single-cell RNA-seq data sets (Park et al., Science, 2020). One included 30,694 cells spanning early thymic development (HSC), and the second contained 76,994 cells from human thymic tissue (Thymus). The right panel shows 26 thymic and hematopoietic cell types that have been defined for each cell by the authors of the scRNA-seq data set. In **Supplementary Table 14** cell type definitions for the abbreviations are provided. (**B**) UMAP plots with individual cell type definitions colored and grouped into six broader categories. (**C**) Heatmap showing ssGSEA for pseudobulks generated for each sample from the single cell reference and using gene sets defined for each cell type. Hierarchical clustering showed a clear separation of developmental stages according to the enrichment scores.

### Supplementary Figure 15

A

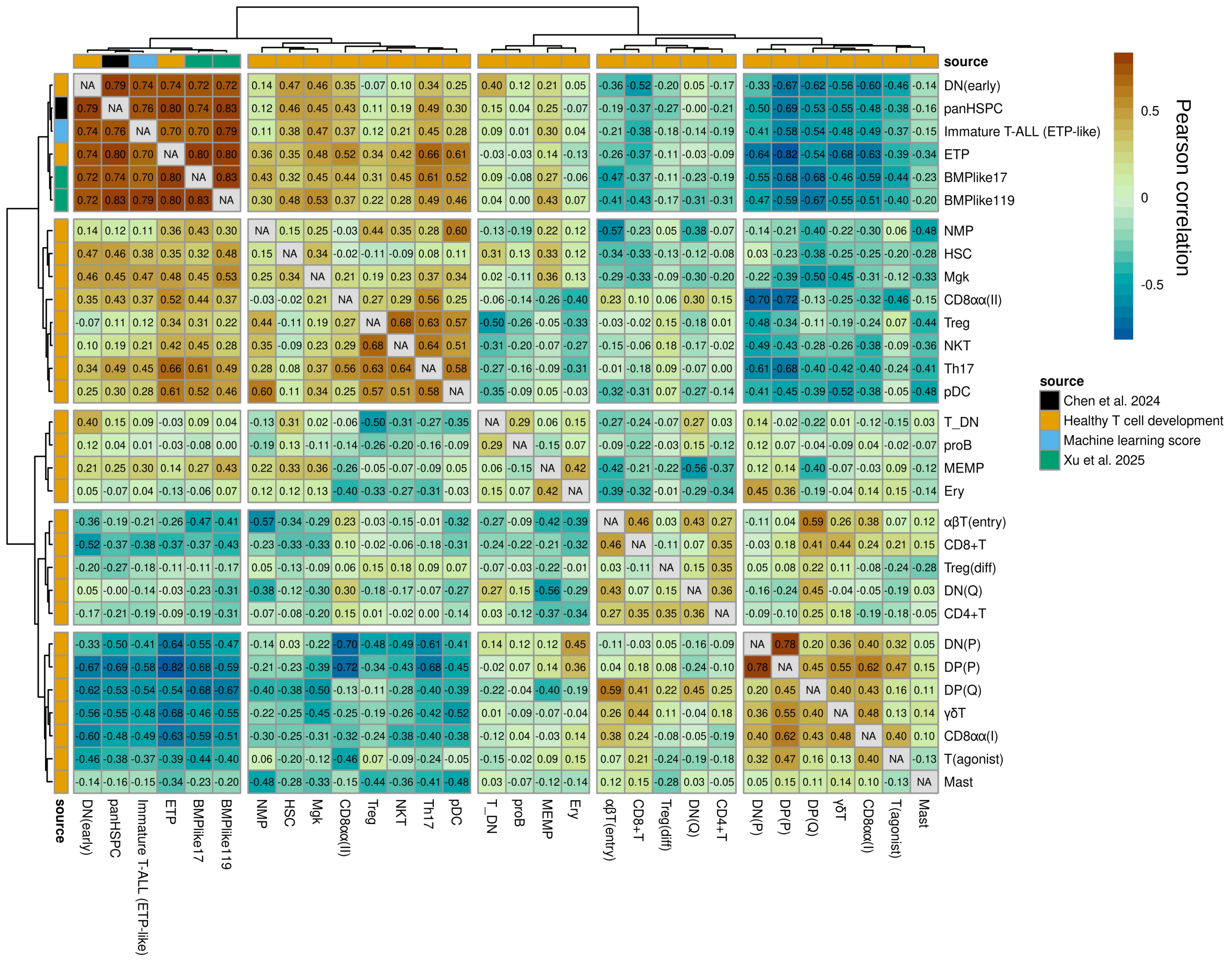


B

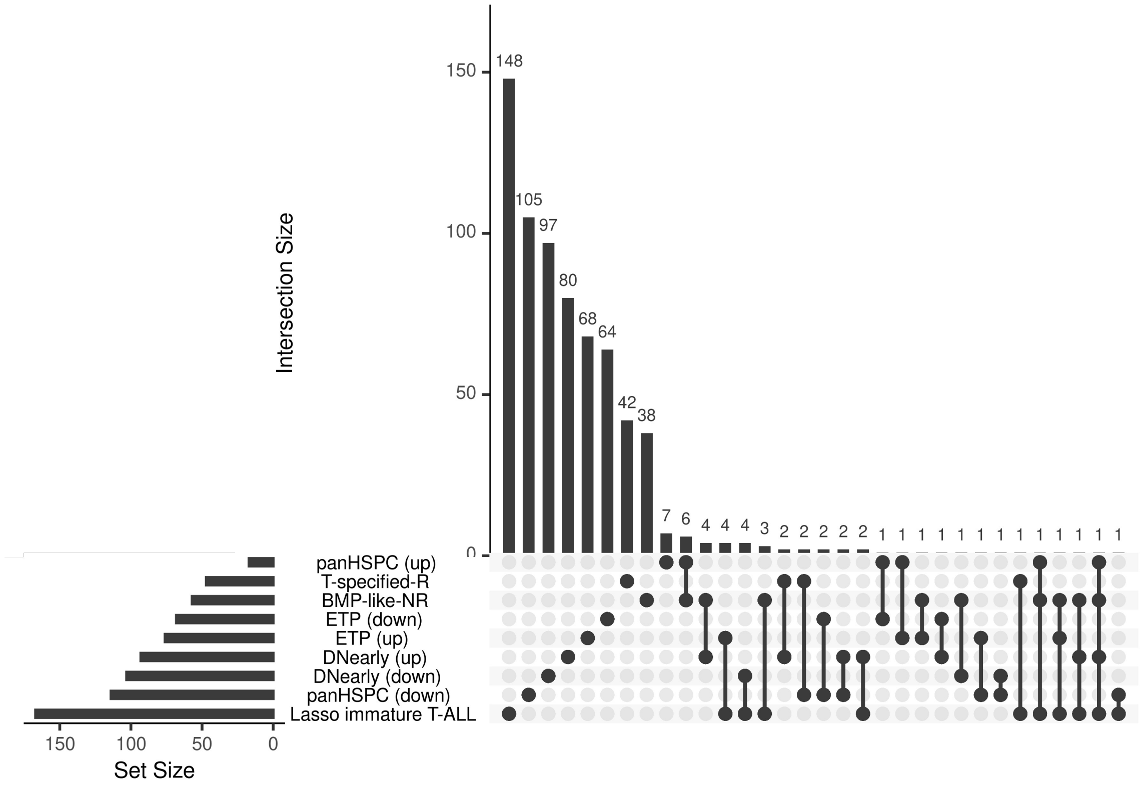


**Supplementary Figure 15. ALLCatchR2 predictions and gene set enrichment analyses of early T cell progenitors correlate with published T-ALL progenitor subpopulation scores.** (**A**) Machine learning prediction scores for immature T-ALL (lightblue) correlated with gene set enrichment to ETP and DN(early) T cell developmental stages, and published scores from single cell T-ALL bone marrow progenitor (BMP)-like subpopulation (Xu et al., Nature Cancer 2025) and hematopoietic stem and progenitor populations in various leukemia (panHSPC, Chen et al., Blood 2025). Values in the heatmap indicate Pearson correlation coefficient. (**B**) Upset plot showing number and intersection between gene sets defined for immature T-ALL (ETP-like), healthy T cell developmental stages and progenitor populations in single cell T-ALL data.
